# AI-m6ARS: Machine learning-driven m6A RNA methylation site discovery with integrated sequence, conservation, and geographical descriptors

**DOI:** 10.1101/2024.06.17.599439

**Authors:** Korawich Uthayopas, Alex G. C. de Sá, David B. Ascher

## Abstract

N6-Methyladenosine (m6A) is a predominant type of human RNA methylation, regulating diverse biochemical processes and being associated with the development of several diseases. Despite its significance, an extensive experimental examination across diverse cellular and transcriptome contexts is still lacking due to time and cost constraints. Computational models have been proposed to prioritise potential m6A methylation sites, although having limited predictive performance due to inadequate characterisation and modelling of m6A sites. This work presents AI-m6ARS, a novel model that utilises integrated sequence, conservation, and geographical descriptive features to predict human m6A methylation sites. The model was trained using the Light Gradient Boosting Machine (LightGBM) algorithm, which was coupled with comprehensive feature selection to improve the data quality. AI-m6RS demonstrates strong predictive capabilities, achieving an impressive area under the receiver operating characteristic curve of 0.87 on cross-validation. Consistent results on unseen transcripts in a blind test highlight the AI-m6ARS generalisability. AI-m6ARS also demonstrates comparable performance to state-of-the-art models, but offers two significant benefits: the model interpretability and the availability of a user-friendly web server. The AI-m6ARS web server offers valuable insights into the distribution of m6A sites within the human genome, thereby facilitating progress in medical applications.

**GRAPHICAL ABSTRACT:** 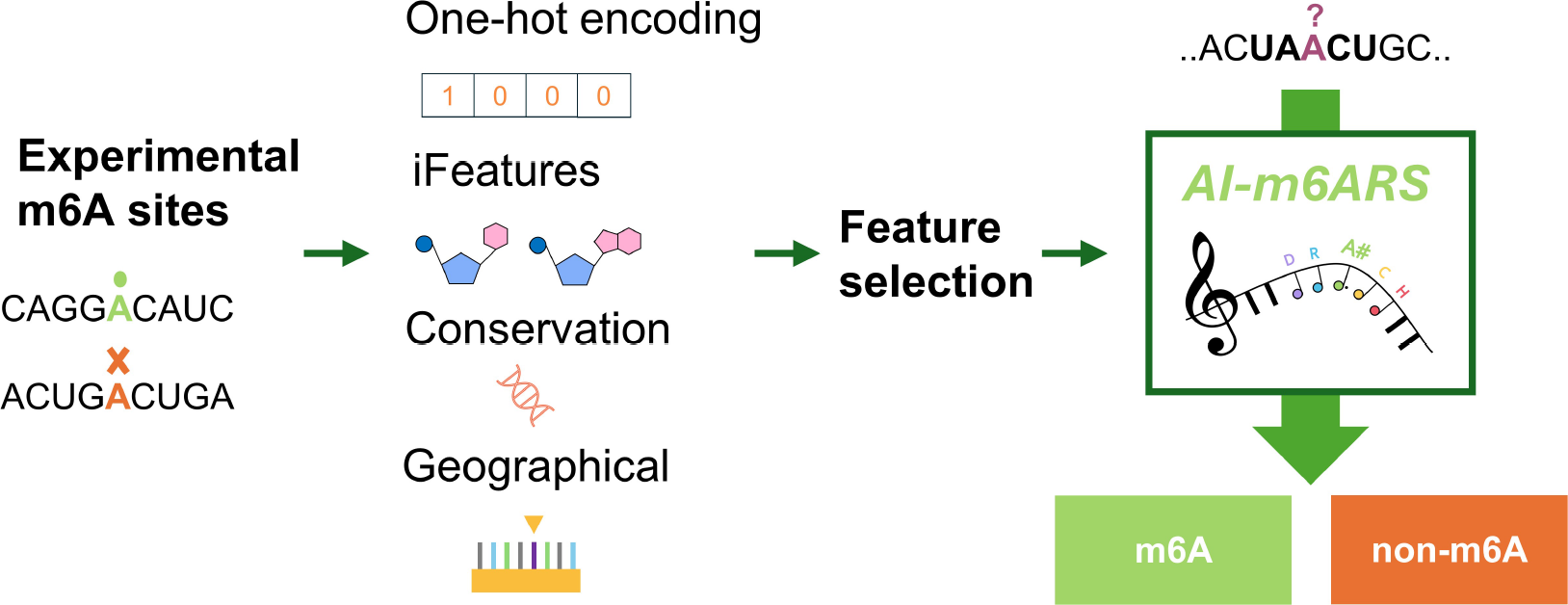

## INTRODUCTION

Gene regulation plays a vital role in controlling gene expression, enabling cells to maintain their optimal functions. Among various gene regulatory mechanisms, post-transcriptional chemical modifications of RNA molecules stand out as an essential component of regulatory systems (1-3). The landscape of RNA modifications is exceptionally diverse, encompassing over 100 identified types in humans to date (1-3). N6-methyladenosine (m6A) emerges as a prevalent and extensively characterised modification within human mRNA (4,5). The reversible nature of m6A modification grants cells the ability to dynamically respond to internal and external stimuli, exerting influence over RNA stability, subcellular localisation, and translational efficiency (6,7).

Due to the properties of the METTL3/METTL4 methyltransferase complex, m6A is primarily deposited within the conserved recognition motif called the DRACH motif (D = A, G, or T; R = A or G; H = A, C, or T) (8,9). Furthermore, m6A was found to cluster predominantly around stop codons (10,11) and away from splice sites (12,13).

Several experiments have conclusively shown a substantial association between m6A methylations and the development of various diseases, such as cancers, neurological disorders, and cardiovascular diseases (4,14-16). Genetic studies known as genome-wide association studies (GWAS) have revealed an array of single nucleotide polymorphisms (SNPs) that interfere with the deposition of m6A, resulting in cellular changes and disease progression (16,17). Understanding the m6A deposition mechanism provides valuable insights into the pathogenesis of diseases, thereby facilitating the investigation of innovative therapeutic and diagnostic approaches.

The rapid progress in transcriptome-wide experimental techniques for accurately identifying modifications at the single-nucleotide resolution has resulted in an accumulation of extensive data regarding the positions of m6A modifications throughout the human transcriptome (18). These methods can be broadly categorised into Antibody-based (18-20) Enzyme-based (8,21,22) Fusion domain-based (22), and Direct nanopore RNA sequencing (DRS) approaches (23,24). The detailed explanation of experimental methodologies employed for the identification of a m6A site has been reviewed in the Supplementary Literature Review.

Antibody-based methods are the main approach for studying m6A modifications, providing a significant amount of data and are commonly used in a number of analyses (18-20). The complex interplay between diverse cellular environments and RNA sequence and structural contexts contribute to specific m6A expression patterns in different cell types. For a complete understanding of the human m6A landscape, an exhaustive examination of the entire human genome in various cell types is required, despite being recognised as a laborious and expensive task. This highlights the necessity for effective computational methods to accurately predict m6A sites in the human transcriptome.

Machine learning models offer several advantages for addressing biological problems, primarily owing to their capability to recognise complex patterns from a large dataset (25). More than ten machine learning models have been proposed in the last decade to predict human m6A methylation sites (26-44). A broad overview of each model is summarised in Table S1. These models have made significant contributions in terms of novel sets of features and algorithms, which resulted in better prediction accuracy. A wide range of algorithms, such as Support Vector Machine (SVM) (26,28,29,33), Random Forest (RF) (27,35,36), Extreme Gradient Boosting (XGBoost) (30,41), and Convolutional neural network (CNN) (32,34,37,40,42,43) were employed.

Despite substantial efforts, the predictive capability of previous models is still limited. It is primarily attributed to the poor characterisation of m6A sites, which relies on a limited variety of feature representations. Expanding feature sets to incorporate additional information shows potential for improving model performance. Significant advancements have been observed in the development of m6A prediction models through the integration of complex sequence-based features, including RNA word embedding (32) and physico-chemical-based one-hot encoding (26,33,35,36). Introducing innovative and extensive sets of sequence features has the capacity to enhance the characterisation of an m6A site sequence and the performance of the model.

Apart from sequence-based features, geographical features have been integrated into multiple studies (33,35,36,41) to characterise the geographic distribution of m6A sites within transcripts, reflecting the preference for m6A deposition for particular regions (10,11). These features include the distance from the exon boundaries and transcript composition (33,35,36,41,42). The integration of a comprehensive and extensive collection of geographical features has yielded notable improvements in model performance, emphasising the crucial significance of the geographical landscape (42).

Prior studies (45,46) have provided compelling evidence suggesting the existence of negative selection within m6A sites (46). Negative or purifying selection is an evolutionary force that maintains physiological functions by eliminating deleterious genetic variations (47,48). Functional regions often exhibit a high degree of conservation, making them useful for identifying functional m6A methylations. Nevertheless, the significance of conservation-based characteristics, which evaluate the level of conservation in m6A sites and neighbouring areas, has been overlooked. Previous models have only utilised two features from the phastCons score (41). Further characterisation could potentially improve the model performance by providing an in-depth analysis of the evolutionary conservation patterns associated with m6A sites.

In addition to concerns about model performance, the accessibility of models poses a significant challenge. The analysis of the 19 models presented in Table S1 reveals that nine models are not accessible (26,28,29,31,32,36-38,41), five models are exclusively available in source code (30,34,39,42,44), two models provide pre-calculated prediction results (33,35), and only three models can be accessed via web servers (27,40,43). Establishing user-friendly web servers is crucial in order to enhance the accessibility of the models. This will enable a wider range of users to explore m6A methylation sites in the human genome.

We introduce AI-m6ARS, an innovative model designed for the Accurate Identification of m6A RNA methylation Sites using machine learning. Addressing the limitations of existing computational methods in predicting m6A RNA methylation sites, AI-m6ARS employs broad approaches to enhance the characterisation of the sequence landscape surrounding an m6A site and its neighbouring region. The feature extraction tool known as iLearn (49,50) has been specifically developed to improve the encoding of physicochemical features derived from DNA, RNA, and protein sequences for solving diverse biological problems (51,52). By employing a combination of 25 sequence encoding methods provided by iLearn along with one-hot encoding, extensive sequence-based features were generated.

Through an in-depth literature review, AI-m6ARS has integrated a comprehensive collection of geographical features and conservation scores. The degree of conservation in m6A and neighbouring sites is represented by two basewise conservation metrics, PhyloP and PhastCons (53,54), which are calculated from multiple sequence alignments of 100 and 30 Vertebrate species.

To improve data quality, Cluster Database at High Identity with Tolerance (CD-HIT) algorithm (55) were employed to reduce redundancy in datasets. The comprehensive feature selection pipeline was implemented to identify an effective and relevant set of features. The AI-m6ARS model, constructed using the LightGBM algorithm (56) and incorporating 175 features, facilitates a comprehensive characterisation of m6A methylation sites, encompassing sequence information, evolutionary conservation, and geographical locations. With these attributes, the AI-m6ARS model accurately predicts m6A sites in the human genome with a single-nucleotide resolution. We believe that these characteristics of AI-m6ARS would greatly contribute to a better understanding of m6A mechanisms in epigenetics research.

## MATERIAL AND METHODS

### AI-m6ARS workflow

The development of AI-m6ARS comprises four steps, as depicted in Figure 1. The validated m6A modification sites were initially obtained from miCLIP experiments (27). Subsequently, the m6A sites were subjected to feature engineering to delineate their characteristics, resulting in a total of 6,396 features. Afterwards, a comprehensive feature selection technique was employed to identify a minimal but effective set of features for training the machine learning model. Thereafter, the AI-m6ARS model was created based on a LightGBM algorithm (56) to identify human m6A methylation sites that are centred within DRACH motifs.

**Figure 1.**
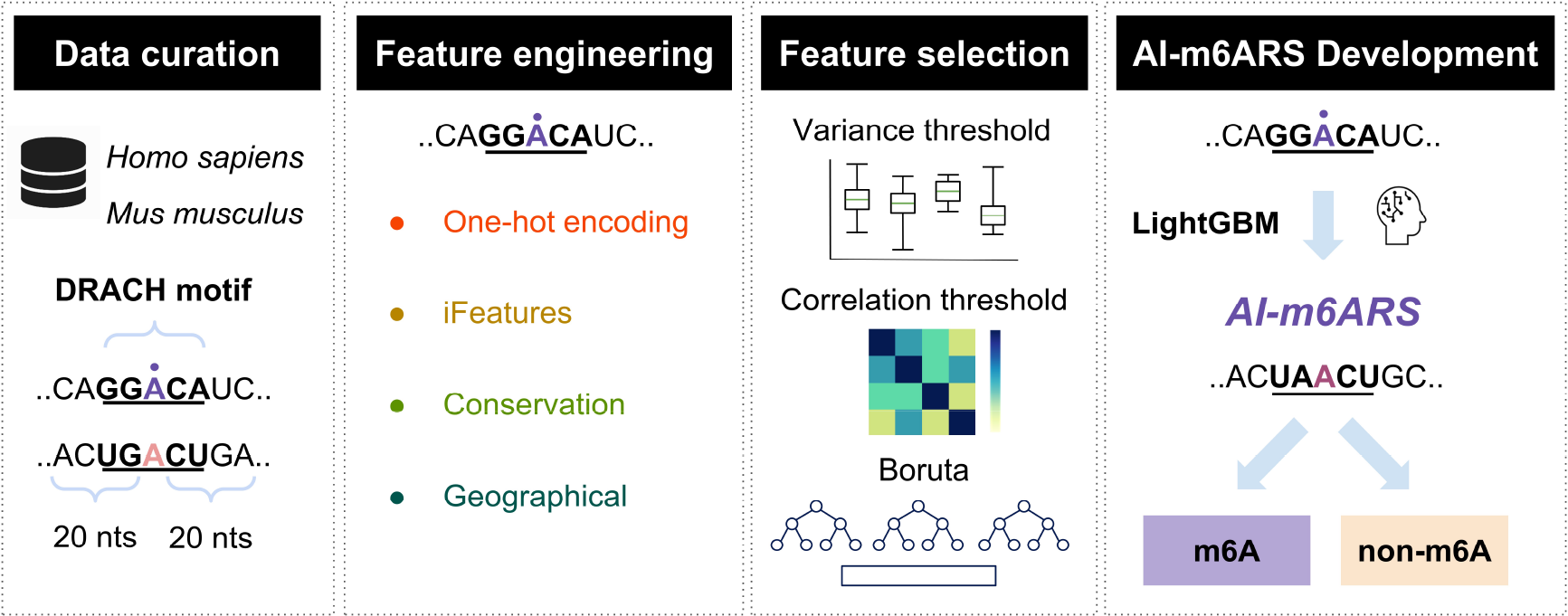
AI-m6ARS workflow: predicting human m6A methylation sites. The development of AI-m6ARS is divided into four steps: (1) Data curation - m6A methylation site within DRACH motifs are validated from multiple experiments, (2) Feature engineering - four sets of features are generated to characterise an m6A methylation site, (3) Feature selection - correlation threshold and Boruta method are employed to select relevant features, (4) AI-m6ARS Development - LightGBM algorithm was employed to predict potential m6A sites in RNAs.

### Data collection and preprocessing

Data collection starts with a curation of human m6A sites within DRACH motifs, a conserved motif for methyltransferase in mammalian m6A. A list of m6A sites was retrieved from miCLIP experiments called the SRAMP dataset (56). The data is then updated using the latest Ensembl version (v109) (57), and the corresponding sequences are obtained. Each datapoint was represented with a 41-nucleotide sequence centred on m6A sites (Figure S1), as similar to previous human m6A predictors (40-44). To improve the specificity of our dataset, only m6A sites that are located within DRACH motifs are included.

Data redundancy presents a significant challenge that can impede the accuracy of the model and introduce biases. To resolve this issue, we adopted a method to eliminate duplicate samples that have identical 41-nucleotide sequences. Out of the duplicate sequences, only one sample was selected at random. Subsequently, the data redundancy issue was further addressed by utilising a sequence similarity metric called CD-HIT. CD-HIT is a renowned technique that was initially created for clustering amino acid and nucleotide sequences (55). This work utilises the CD-HIT algorithm with a threshold of 0.8 to remove sequences that are statistically similar. These methods effectively reduced the redundancy in our dataset, resulting in improved performance of the machine learning model.

As a result, a training dataset comprising 8,811 confirmed m6A sites and 82,187 non-m6A sites was derived from 2,276 transcripts. These sites were classified as positive and negative samples, respectively. For evaluation, an independent blind test set was obtained from SRAMP and subjected to identical preprocessing procedures. The blind test consists of 2,189 positive samples and 20,204 negative samples, which covers 553 transcripts. Notably, the transcripts in the blind test set are completely different from those in the training set to ensure an unbiased evaluation.

Additionally, m6A sites specific to the mouse (Mus musculus) genome were obtained from the SRAMP dataset. It results in a training set of 13,223 positive and 116,284 negative samples from 4,856 transcripts. The blind test set comprises 4,359 positive and 39,164 negative samples from 1,208 non-overlapping transcripts.

### Feature engineering

A comprehensive set of 6,396 features is created to characterise a m6A methylation site. The features include 164 one-hot encoding features, 4,591 iFeatures, 36 conservation scores, and 1,605 geographical features. The following features are briefly summarised below. An in-depth description of a set of features is presented in Table S2.

#### One-hot encoding (164 features)

One-hot encoding involves encoding four types of nucleobases into binary vectors, resulting in 164 distinct features representing 41 nucleotide sequences centred on m6A methylation sites.

#### iFeatures (4,591 features)

iFeatures utilised in this study are a collection of sequence-derived features obtained from the iLearn Python toolkit (51,52). iLearn was developed to generate a comprehensive array of features from DNA, RNA, and protein sequences that can accurately represent their physical and chemical properties using multiple methods (51,52). Its versatility has been demonstrated in diverse real-world applications, such as the identification of promoters (58) and pseudouridine sites (59).

The present study utilised a total of 25 unique RNA sequence encoding schemes available in iLearn in order to investigate the physicochemical characteristics of the m6A sequences. The schemes include Kmer type 1, Mismatch, Subsequence, Enhanced nucleic acid composition (ENAC), Accumulated nucleotide frequency (ANF), binary, Position-specific of two nucleotides (PS2), Position-specific of three nucleotides (PS3), Composition of k-spaced nucleic acid pairs (CKSNAP) type 1, Nucleotide chemical property (NCP), Adaptive skip dinucleotide composition (ASDC), Dinucleotide binary encoding (DBE), Local position-specific dinucleotide frequency (LPDF), Dinucleotide physicochemical properties (DPCP) type 2, Z curve 9bit, Z curve 12bit, Z curve 36bit, Normalised moreau-broto autocorrelation (NMBroto), Moran, Geary, dinucleotide-based cross covariance (DCC), dinucleotide-based auto-cross covariance (DACC), pseudo k-tuple nucleotide composition (PseKNC), parallel correlation pseudo dinucleotide composition (PC-PseDNC), and series correlation pseudo dinucleotide composition (SC-PseDNC).

#### Conservation scores (36 features)

Conservation scores represent a degree of conservation within the m6A methylation site, DRACH motif, and adjacent regions. A high degree of conservation has been shown to be a distinguishing feature of functional m6A sites in previous research (45,46). m6A sites subject to significant evolutionary constraints demonstrate higher levels of methylation. These sites are hypothesised to be optimised for m6A writer binding and subsequent methylation (46) Two basewise conservation metrics, PhyloP (53) and PhastCons scores (54), have been introduced to delineate the conservation profile.

PhyloP evaluates the evolutionary conservation of specific nucleotide positions by comparing them with other related species (53). Conservation at the site is indicated by positive scores, while rapid evolution of the site is indicated by negative scores. PhastCons, on the other hand, is specifically designed to identify conserved regions, offering a broader perspective on conservation (54). The pre-computed scores, derived from multiple sequence alignments involving 100 and 30 vertebrate species, were obtained from the UCSC Genome Browser (60). AI-m6ARS utilise PhyloP and PhastCons to characterise three regions: the m6A site, the DRACH motif, and the neighbouring regions. Mean values were selected to represent the conservation level of these regions.

#### Geographical features (1,605 features)

Geographical features encode the spatial information relevant to the m6A methylation site. Previous work, Geography Plus Sequences (GepSe) (42), has presented a comprehensive set of features called Geographical representation of transcript as vectors (Geo2Vec), which consists of landmarkTX, gridTX, and chunkTX. Based on Geo2Vec, we have generated a comprehensive set of geographical features encompassing three types of information - 15 distance-based features, 1080 fragment-based features, and 510 region-based features (Figure S2).

Distance-based features capture essential positional information of m6A sites within transcripts, including distances to geographical landmarks such as the start and end of the transcript, exons, splice sites, 5’-UTR, and 3’-UTR. A total of 15 features provide an overview of the relative positioning of m6A sites in relation to the landmark.

Fragment-based features were generated to enhance the resolution of geographical features. The transcript was partitioned into a specific number of fragments, with each fragment being represented by nine different features (Table S2). These features indicate the composition of specific region types within the fragment, calculated by dividing the number of nucleotides in those regions by the length of the fragment. The types of specific regions include exons, 5’-UTR, coding sequences (CDS), introns, and 3’-UTR. Varying the number of fragments (8, 16, 32, and 64) enables the generation of features at different levels of detail. It allows the model to recognise the local information of each fragment and its relative position within the transcripts.

Region-based features work at the regional level, capturing information such as the width, composition, and relative order of each region. The transcripts were split into a number of regions according to their region types, including exon, 5’-UTR, CDS, intron, and 3’-UTR. 10 features, as shown in Table S2, encode the presence of the m6A site, region length, and region type for each region. In consideration of the diverse numbers of regions in different transcripts, the technique of zero-padding was utilised for simpler transcripts, while trimming was employed for more complex transcripts. The m6A site is located at the centre of the feature matrix during a feature generation with a width of 1.

AI-m6ARS has made notable improvements from previous studies by expanding a set of geographical features. A distance to the splice site was incorporated, which acknowledges the m6A preferential deposition outside splice site regions (13). Fragment-based features employ a range of fragments instead of a fixed number, providing more detailed information at diverse resolutions. In region-based features, a proportion of region length to the entire transcript is computed to standardise across transcripts with different lengths.

### Feature preprocessing and selection

A feature selection process was performed on the initial set of 6,396 features to improve predictive performance, decrease model complexity, and reduce computational workload (25,61). In order to determine the most effective feature selection pipeline for our dataset, we conducted a thorough evaluation considering the wide range of available feature selection methods and data scaling options. We employed a validation set consisting of 20% randomly chosen samples from the training dataset. A blind test and 5-fold cross-validation were conducted using a Light Gradient Boosting Machine (LightGBM) model, which has been determined to be the most accurate when trained with a complete set of features (Table S2).

Examining 126 unique combinations, we evaluated seven techniques for scaling data [Hands-On Machine Learning with Scikit-Learn, Keras, and TensorFlow, 2nd Edition] (No scaling, StandardScaler, MinMaxScaler, MaxAbsScaler, RobustScaler, QuantileTranformer, PowerTranformer), three options for Variance Threshold (62,63) (No threshold, 0.95, 0.99), three options for Correlation Threshold (63,64) (No threshold, 0.95, 0.99), and two options for Boruta (65) (executed, not executed). Supplementary Materials and Methods contain comprehensive information on the employed data scaling techniques, variance threshold, correlation threshold, and the application of Boruta.

The performances of the models across all combinations were evaluated and ranked using Matthew’s correlation coefficient (MCC) (25,66), as provided in Table S3. MCC is a reliable metric for evaluating the predictive performance of binary classification models. By considering true positives, true negatives, false positives, and false negatives in its calculation, MCC yields high scores only when the model performs exceptionally well in all metrics, preventing the propagation of over-optimistic results (67).

### Model training, optimisation, validation

A range of supervised machine learning algorithms was independently assessed, including Light Gradient Boosting Machine (LightGBM) (56), Gradient Boosting (GB) (68) Explainable Boosting Machine (EBM) (69), Extreme Gradient Boosting (XGBoost) (70), Adaptive Boosting (AdaBoost) (71), Random Forest (RF) (72), Extra Trees (73), Multilayer perceptron (MLP) (74), K-Nearest Neighbors (KNN) (75), and LogisticRegression (76). Each algorithm has its strengths and weaknesses, and the choice of the best algorithm depends on various factors such as dataset size, sparsity, feature types, and computational resources.

LightGBM exhibited superior performance compared to other algorithms in training AI-m6ARS using the complete feature set of 6,396 features (Table S2). LightGBM, developed by Microsoft, is a gradient boosting framework that utilises a leaf-wise growth strategy (56). The advantages of this algorithm include high training efficiency, reduced memory consumption, scalability for handling extensive datasets, and improved predictive capabilities. As a result, the LightGBM algorithm was selected for the training of AI-m6ARS with 175 selected features. Hyperparameter optimisation was subsequently performed to fine-tune the algorithm’s parameters, such as a learning rate, which controls the learning process to achieve peak predictive performances and maximise model generalisability on a given training set. We employed Bayesian optimisation, offered by the scikit-optimize library, to effectively tune the hyperparameters of the LightGBM model.

The AI-m6ARS model was developed using a set of human m6A sites and the LightGBM algorithm. Evaluating the performance of the model across various organisms provides valuable insights into its capability to adapt in diverse contexts, such as genomes and cellular environments. A mouse-specific AI-m6ARS model was developed and trained independently using mouse m6A sites obtained from the SRAMP dataset (27). An in-depth examination of the model trained using a combination of data from different species provides valuable insights into the potential improvements that can be achieved by integrating information from various species. This approach helps to overcome the limitations imposed by the low data availability. The third AI-m6ARS model was trained using a dataset comprising m6A sites from both humans and mice in SRAMP dataset. Each model, including human, mouse, and human-mouse, was subjected to an independent AI-m6ARS pipeline. Notably, the exclusion of conservation features from the pipeline during training with the mouse and human-mouse was primarily due to the unavailability of employed conservation scores in mice.

Internal cross-validation approaches, as well as external validation approaches with independent test sets, were performed to evaluate the performance of AI-m6ARS using different metrics including the Area Under the Receiver Operating Characteristic Curve (AUC), Balanced accuracy (bACC), F1-score (F1), MCC, Precision, Recall, and Specificity (25,66). Detailed information about the performance metrics employed is available in Supplementary Materials and Methods.

### Model interpretation

Model interpretation is the fundamental procedure of understanding the decision-making mechanisms of a machine learning algorithm, specifically how a model generates predictions (77). It has the ability to detect and reduce biases present in machine learning models and provides a way to improve model performance and generalisability.

The study utilises SHapley Addictive exPlanations (SHAP) analysis to offer an insight into the model’s output (78). SHAP analysis is a widely utilised technique in diverse academic fields, serving as a fundamental approach for the interpretation of machine learning models (79,80). Based on game theory principles, SHAP values are computed to evaluate the impact of each feature on prediction results. Features that exhibit positive SHAP values are indicative of their positive contribution to predictions, suggesting their association with functional m6A sites. In contrast, negative values indicate features associated with non-functional m6A sites or negative samples. Furthermore, the magnitude of SHAP values serves to clarify the extent of influence on the model’s prediction.

## RESULTS

### Feature selection

The performances of various combinations of data scaling and feature selection methods were evaluated using MCC (Table S3). The most effective pipeline comprises a variance threshold of 0.95, a correlation threshold of 0.95, and the Boruta algorithm. The pipeline yields a concise yet effective set of 175 features, comprising 80 iFeatures, 6 conservation scores, and 89 geographical features (Table S4). All feature types are included, with the exception of one-hot encoding. This observation indicates that the utilisation of a collection of complex sequence encoding schemes, such as those provided by iLearn (51,52), might be more advantageous in model training compared to one-hot encoding. Additionally, it highlights the importance of incorporating sequence, geographical, and conservation information to achieve an effective representation of m6A sites.

Examining the selected iFeatures, out of 25 sequence encoding methods, only four types were selected. These include 38 Mismatch, 4 Subsequence, 29 CKSNAP, and 9 DPCP type 2 (Table S4). Mismatch and Subsequence, variants of Kmer that allow mismatches (81) and non-contiguous matches (82), consider RNA sequence as words, providing advantages over unselected standard Kmer in accurately representing the sequence contexts of m6A sites.

CKSNAP calculates the frequency of nucleic acid pairs separated by any k nucleic acid. On the other hand, DPCP type 2 encodes the physicochemical properties of consecutive RNA dinucleotides (83), specifically the angular characteristics of base pairs located at positions 10, 11, 30, 37, and 38 within a 41-nt region (Table S2).

The six selected conservation scores that have been selected are exclusively obtained from the 100-way PhyloP (Table S4). The scores include conservation scores from m6A sites and D, R, C, and H positions in DRACH motifs. A mean value of conservation scores is taken to represent the conservation of a 41-nt region. The exclusion of 30-way PhyloP, 100-way PhastCons, and 30-way PhastCons from feature selection suggests that the 100-way PhyloP alone is sufficient for characterising the evolutionary conservation patterns of m6A and neighbouring sites. Possible explanations may include the computation of PhyloP, which evaluates conservation at specific nucleotide positions while disregarding the influence of neighbouring sites (53), and the enhanced precision of conservation levels provided by alignments of 100 vertebrates as opposed to 30 vertebrates.

The geographical features consist of seven distance-based features, one fragment-based feature, and 81 region-based features (Table S4). Distance-based features are utilised to depict the distances to important geographical landmarks, including transcript boundaries, 5’-UTR, 3’-UTR, exons, and splice sites (Figure S2). A selected fragment-based feature denotes the length of the last fragment when a transcript is divided into 64 fragments (Figure S2). Region-based features, which encode the length of each region and the proportion of region length to the entire transcript (-log_2_), capture the landscape of region lengths. Although the feature selection does not directly select region-type features within region-based features, the model can still inherently identify the region by considering its size and relative position.

### Performance of AI-m6ARS

Different cross-validation approaches were employed to evaluate the predictive performance of AI-m6ARS for predicting m6A methylation sites in human transcripts. On 5-fold cross-validation, AI-m6ARS achieved AUC, bACC, F1, and MCC values of 0.872, 0.746, 0.477, and 0.419, respectively. In addition, AI-m6ARS yielded comparable results on 10- and 20-fold cross-validation approaches, indicating its ability to accurately predict m6A methylation sites in the human genome (Table 1 and Figure 2).

**Figure 2.**
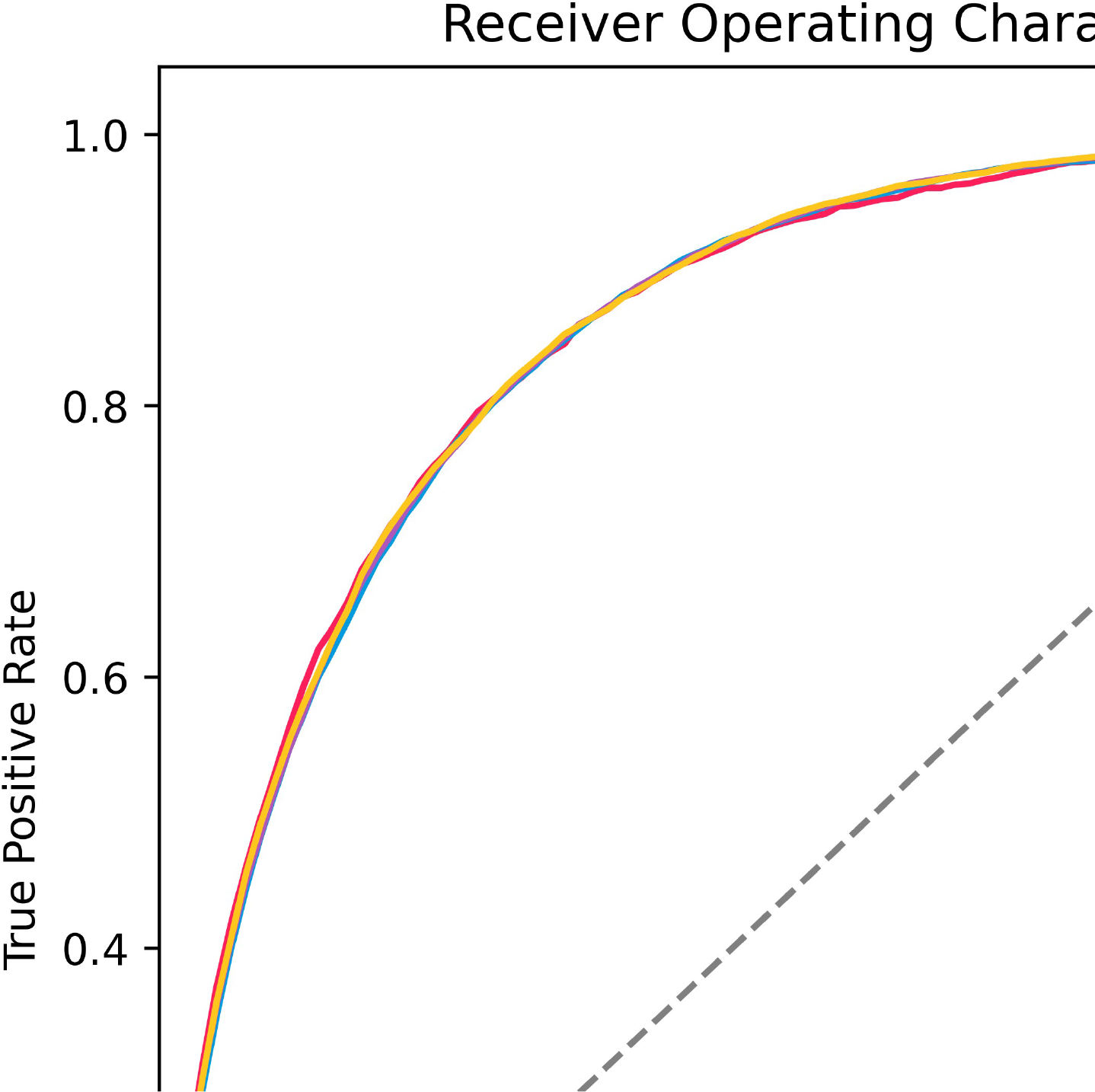
Receiver Operating Characteristic (ROC) Curve Analysis of AI-m6ARS. The AI-m6ARS model demonstrates high predictive performance in predicting m6A methylation sites in the human genome, achieving AUC values of 0.861, 0.872, 0.874, and 0.875 in a blind test, 5-fold, 10-fold, and 20-fold cross-validation, respectively.

**Table 1.** AI-m6ARS performance evaluation on 5-fold, 10-fold, and 20-fold cross-validation, and a blind test.

| Methods | AUC | bACC | F1 | MCC | Precision | Recall | Specificity |
| --- | --- | --- | --- | --- | --- | --- | --- |
| 5-fold cross-validation | 0.863 ± 0.004 | 0.746 ± 0.006 | 0.453 ± 0.008 | 0.395 ± 0.009 | 0.361 ± 0.006 | 0.607 ± 0.012 | 0.885 ± 0.002 |
| 10-fold cross-validation | 0.865 ± 0.005 | 0.754 ± 0.010 | 0.454 ± 0.013 | 0.398 ± 0.015 | 0.355 ± 0.011 | 0.630 ± 0.020 | 0.878 ± 0.005 |
| 20-fold<br>cross-validation | 0.866 ±<br>0.010 | 0.756 ±<br>0.015 | 0.453 ±<br>0.020 | 0.398 ±<br>0.024 | 0.352 ±<br>0.017 | 0.636 ±<br>0.028 | 0.875 ±<br>0.007 |
| Blind test | 0.865 | 0.756 | 0.465 | 0.408 | 0.369 | 0.628 | 0.883 |

Additionally, we conducted an assessment across an independent blind test to estimate AI-m6ARS generalisability in newly unseen transcripts. The model has achieved AUC, bACC, F1, and MCC scores of 0.865, 0.756, 0.465, and 0.408 respectively. These results are consistent with the predictive results from cross-validation procedures (Table 1 and Figure 2).

To assess the generalisation capabilities of AI-m6ARS in contrast to previous state-of-the-art models, GepSe (42), SRAMP (27), DL-m6A (43), and TS-m6A-DL (40), we utilise an independent blind test from SRAMP (27). After removing data points that are not supported by GepSe, the remaining dataset was used as input for the other models. TS-m6A-DL consists of three models specifically designed for each human tissue (40). All models were used to generate prediction results. Our analysis demonstrates that AI-m6ARS surpasses other state-of-the-art models. Comparatively, AI-m6ARS exhibits a slightly improved overall performance in terms of MCC and bACC when compared to GepSe (Table 2).

**Table 2.** Performance comparison of state-of-the-art models on the human SRAMP blind test.

| **Models** | **AUC** | **bACC** | **F1** | **MCC** | **Precision** | **Recall** | **Specificity** |
| --- | --- | --- | --- | --- | --- | --- | --- |
| AI-m6ARS | 0.867 | 0.758 | 0.471 | 0.414 | 0.376 | 0.630 | 0.886 |
| GepSe | 0.868 | 0.735 | 0.471 | 0.409 | 0.407 | 0.559 | 0.911 |
| SRAMP | N/A* | 0.707 | 0.396 | 0.326 | 0.310 | 0.549 | 0.866 |
| DL-m6A<br>(threshold = 0.5) | N/A* | 0.517 | 0.178 | 0.021 | 0.103 | 0.640 | 0.394 |
| TS-m6A-DL kidney<br>(threshold = 0.5) | N/A* | 0.501 | 0.169 | 0.001 | 0.099 | 0.582 | 0.420 |
| TS-m6A-DL brain<br>(threshold = 0.5) | N/A* | 0.498 | 0.167 | -0.003 | 0.098 | 0.561 | 0.434 |
| TS-m6A-DL liver | N/A* | 0.498 | 0.169 | -0.003 | 0.098 | 0.614 | 0.382 |
| (threshold = 0.5) |  |  |  |  |  |  |  |

\*The AUC values are unavailable due to the absence of probability scores in the results obtained from the respective web servers.

The performance of AI-m6ARS when training in other species was evaluated using a curated dataset of m6A sites obtained from the mouse genome, retrieved from the SRAMP dataset (27). The dataset went through the same AI-m6ARS pipeline, resulting in the m6A site predicting model trained from a mouse genome. The findings indicate a great performance of AI-m6ARS on mouse data, as demonstrated by AUC, bACC, F1, and MCC values of 0.860, 0.747, 0.442, and 0.382, respectively (Table 3).

**Table 3.** Cross-species analysis of m6A site prediction models across human and mouse SRAMP datasets.

| Train set | AUC | bACC | F1 | MCC | Precision | Recall | Specificity |
| --- | --- | --- | --- | --- | --- | --- | --- |
| **Test set: Human** |  |  |  |  |  |  |  |
| Human | 0.861 | 0.747 | 0.460 | 0.401 | 0.371 | 0.606 | 0.889 |
| Mouse | 0.862 | 0.736 | 0.468 | 0.407 | 0.401 | 0.563 | 0.909 |
| Combined Human and Mouse | 0.871 | 0.763 | 0.470 | 0.415 | 0.369 | 0.646 | 0.881 |
| **Test set: Mouse** |  |  |  |  |  |  |  |
| Human | 0.845 | 0.744 | 0.417 | 0.358 | 0.307 | 0.652 | 0.836 |
| Mouse | 0.860 | 0.747 | 0.442 | 0.382 | 0.341 | 0.629 | 0.865 |
| Combined Human and Mouse | 0.861 | 0.766 | 0.435 | 0.384 | 0.315 | 0.701 | 0.831 |

To examine the cross-species applicability of the AI-m6ARS pipeline, we conducted tests on the model that was trained using human data on mouse data, and vice versa. Remarkably, the models accurately predict m6A sites across different species, albeit with a slight performance decrease when training with humans and testing on mice. Notably, the model trained on mouse data and evaluated on human data exhibited superior performance than a human model in terms of F1 and MCC, reaching values of 0.468 and 0.407, compared to the human model’s values of 0.460 and 0.401, respectively (Table 3). Two potential factors might contribute to this result. The generalisability of the model might still be good from the presence of shared m6A signatures across closely related species. Furthermore, the higher number of mouse dataset, nearly twice as large as humans, might benefit the model training.

In order to investigate the potential benefits of incorporating human and mouse data into the AI-m6ARS pipeline, a unified model was trained using a combined dataset. Both humans and mice demonstrated a significant enhancement in the overall predictive accuracy for m6A sites, as indicated by AUC, bACC, F1, and MCC. When evaluating in human data the model built in both human and mouse data, these metrics increased from the human-only model’s values of 0.861, 0.736, 0.468, and 0.401 to 0.871, 0.763, 0.470, and 0.415, respectively (Table 3). On the other hand, the predictions made in the mouse dataset demonstrate slight but noticeable improvements in AUC, bACC, and MCC, progressing from 0.860, 0.747, and 0.382 to 0.861, 0.766, and 0.384, respectively (Table 3). These findings reveal that combining experimental data from both mice and humans enhances the model’s predictive ability in both mammals.

### Model interpretation

In order to conduct a comprehensive examination of the machine learning model, we employ SHAP analysis for model interpretation (78). Figure 3 displays the SHAP values, which highlight the top 20 features ranked in descending order according to their influence on model predictions. The features consist of 11 iFeatures, 8 geographical features, and 1 conservation score. This indicates that all types of features play a role in the decision-making process inside the model, showing that integrating both sequence, geographical, and conservation contexts is important.

**Figure 3.**
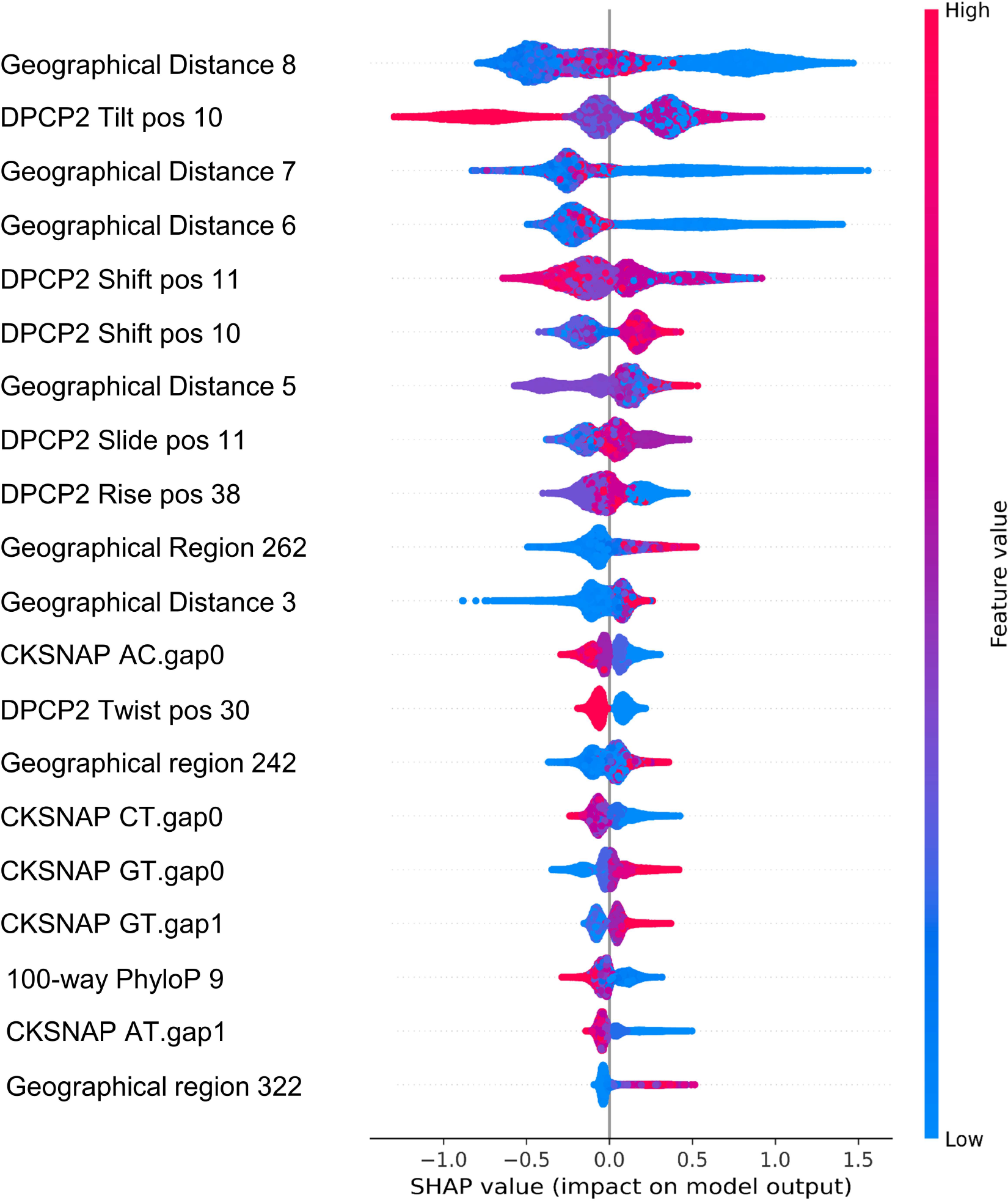
The SHAP analysis of AI-m6ARS. SHAP values were calculated for each individual feature. The feature importance is determined based on their impact on model predictions. Each dot represents a m6A methylation site and is assigned a colour according to its feature value. Points with high values are depicted in a red colour, whereas points with low values are depicted in a blue colour. The top 20 most important features are shown.

The most important feature is the 8^th^ distance-based geographical feature, which reflects the proximity to the nearest splice site. This finding aligns with a previous study that highlighted the significant role of splice-site proximity in m6A topologies and the exclusion of m6A sites through the exon junction complex (13). Additional geographic features were ranked at positions 3, 4, 7, 11, 14, and 20. The majority of them are distance-based, capturing positional information in relation to geographical landmarks, including the end of transcripts, the start of 3’-UTR, and the exon boundaries. In addition, the remaining region-based geographical features (262^nd^, 242^nd^, and 322^nd^) indicate the length of the regions that are next to the target sites, as well as the 8th region on the right side.

The most significant iFeatures are a group of six DPCP type 2 features that encode the physicochemical properties of RNA sequences. Following these, there are five CKSNAP features that represent the nucleotide composition (83). Out of the top 20 features, the only one concerning conservation is the 100-way PhyloP. This score provides an overview of the degree of conservation across the entire 41-nucleotide segment, rather than specifically focusing on the DRACH motif or methylated site.

A comprehensive analysis was performed to examine the contribution of different feature sets to the overall model performance and to evaluate the effectiveness of each feature set to represent m6A sites. Each model was constructed for each set of features, iFeature, conservation scores, and geographical features, with a hyperparameter optimization to ensure optimal performance for each. The evaluation revealed that employing either iFeatures or Geographical features in isolation yielded commendable results, with AUC, bACC, F1, and MCC values surpassing 0.750, 0.650, 0.300, and 0.230, respectively (Table 4). The incorporation of both feature sets demonstrated a significant improvement in performance (Table 4), with AUC, bACC, F1, and MCC reaching approximately 0.860, 0.740, 0.450, and 0.390. The integration of iFeatures and Geographical features is identified as a crucial factor in achieving excellent model performance, emphasising the importance of feature integration for optimal predictive accuracy.

**Table 4.** The performance of the AI-m6ARS Model trained with an individual feature set: iFeatures, Conservation scores, and Geographical features.

| Methods | AUC | bACC | F1 | MCC | Precision | Recall | Specificity |
| --- | --- | --- | --- | --- | --- | --- | --- |
| **iFeatures** |  |  |  |  |  |  |  |
| 5-fold cross-validation | 0.749 ± 0.003 | 0.664 ± 0.006 | 0.320 ± 0.006 | 0.238 ± 0.007 | 0.234 ± 0.005 | 0.504 ± 0.012 | 0.823 ± 0.002 |
| Blind test | 0.758 | 0.673 | 0.330 | 0.251 | 0.240 | 0.528 | 0.819 |
| **Conservation scores** |  |  |  |  |  |  |  |
| 5-fold cross-validation | 0.580 ± 0.004 | 0.557 ± 0.004 | 0.195 ± 0.004 | 0.068 ± 0.005 | 0.120 ± 0.003 | 0.529 ± 0.010 | 0.584 ± 0.009 |
| Blind test | 0.575 | 0.547 | 0.191 | 0.057 | 0.117 | 0.513 | 0.582 |
| **Geographical features** |  |  |  |  |  |  |  |
| 5-fold cross-validation | 0.804 ± 0.005 | 0.720 ± 0.006 | 0.347 ± 0.005 | 0.289 ± 0.007 | 0.232 ± 0.003 | 0.681 ± 0.016 | 0.760 ± 0.004 |
| Blind test | 0.775 | 0.684 | 0.325 | 0.251 | 0.225 | 0.589 | 0.780 |
| **iFeatures + Conservation scores** |  |  |  |  |  |  |  |
| 5-fold cross-validation | 0.756 ± 0.003 | 0.661 ± 0.005 | 0.330 ± 0.005 | 0.249 ± 0.007 | 0.255 ± 0.004 | 0.466 ± 0.010 | 0.855 ± 0.002 |
| Blind test | 0.762 | 0.669 | 0.340 | 0.260 | 0.262 | 0.486 | 0.852 |
| **iFeatures + Geographical features** |  |  |  |  |  |  |  |
| 5-fold cross-validation | 0.860 ±<br>0.003 | 0.737 ±<br>0.005 | 0.450 ±<br>0.009 | 0.389 ±<br>0.010 | 0.366 ±<br>0.010 | 0.583 ±<br>0.009 | 0.892 ±<br>0.004 |
| Blind test | 0.858 | 0.741 | 0.458 | 0.397 | 0.375 | 0.589 | 0.893 |
| **Conservation scores + Geographical features** |  |  |  |  |  |  |  |
| 5-fold cross-validation | 0.804 ±<br>0.003 | 0.729 ±<br>0.003 | 0.342 ±<br>0.004 | 0.292 ±<br>0.004 | 0.224 ±<br>0.003 | 0.730 ±<br>0.011 | 0.730 ±<br>0.006 |
| Blind test | 0.785 | 0.708 | 0.31 | 0.269 | 0.219 | 0.676 | 0.739 |

In contrast, the model that was trained solely on conservation scores demonstrated a limited predictive capability, as indicated by an AUC value of only 0.580 (Table 4). The evaluation of the models that integrated conservation scores with iFeatures and with Geographical features revealed a slight improvement in overall performance (Table 4). These results suggest that the conservation scores may exert only a minor influence on the prediction computation in AI-m6ARS, in comparison to iFeatures and geographical features. This aligns with the evidence from the SHAP analysis, which identified only one conservation feature among the top 20 most important features (Figure 3).

Potential causes of a limited impact can be attributed to the subtle differences in conservation patterns observed between functional and non-functional m6A sites, which may pose a challenge to the model’s capability to distinguish between them. A previous genomics study has provided evidence suggesting that only a small proportion of m6A sites in mammals experience negative selection (45). Furthermore, it has been observed that recently gained m6A sites experience positive selection, as indicated by SNP genotype data (46). The diversity in selection pressures complicates the model’s characterisation of conservation levels. Nevertheless, given their significant contribution toward the improvement of model performance (Table 4), we have decided to incorporate conservation scores into the AI-m6ARS pipeline.

### Online web interface

AI-m6ARS can be accessed through a user-friendly web server accessible at https://biosig.lab.uq.edu.au/aim6ars/. Figure 4 displays the Prediction and Results pages of AI-m6ARS. To start the analysis, users must choose the input mode between Transcript Scanning and Transcript Coordinate, as shown in Figure 4A. AI-m6ARS generates accurate predictions for each DRACH motif present in the input transcripts. On the other hand, the latter mode limits predictions to particular locations within the input transcripts. It is recommended to use the Ensembl Transcript (ENST) format as the input. The AI-m6ARS algorithm provides predictions in a tabular format (as shown in Figure 4B), presenting the number and specific positions of potential m6A sites for each transcript. The Detail page in Figure S3 provides comprehensive information, including transcript details, predicted m6A sites with corresponding probabilities, DRACH motif sequences, and a distribution graph of m6A modification sites.

**Figure 4.**
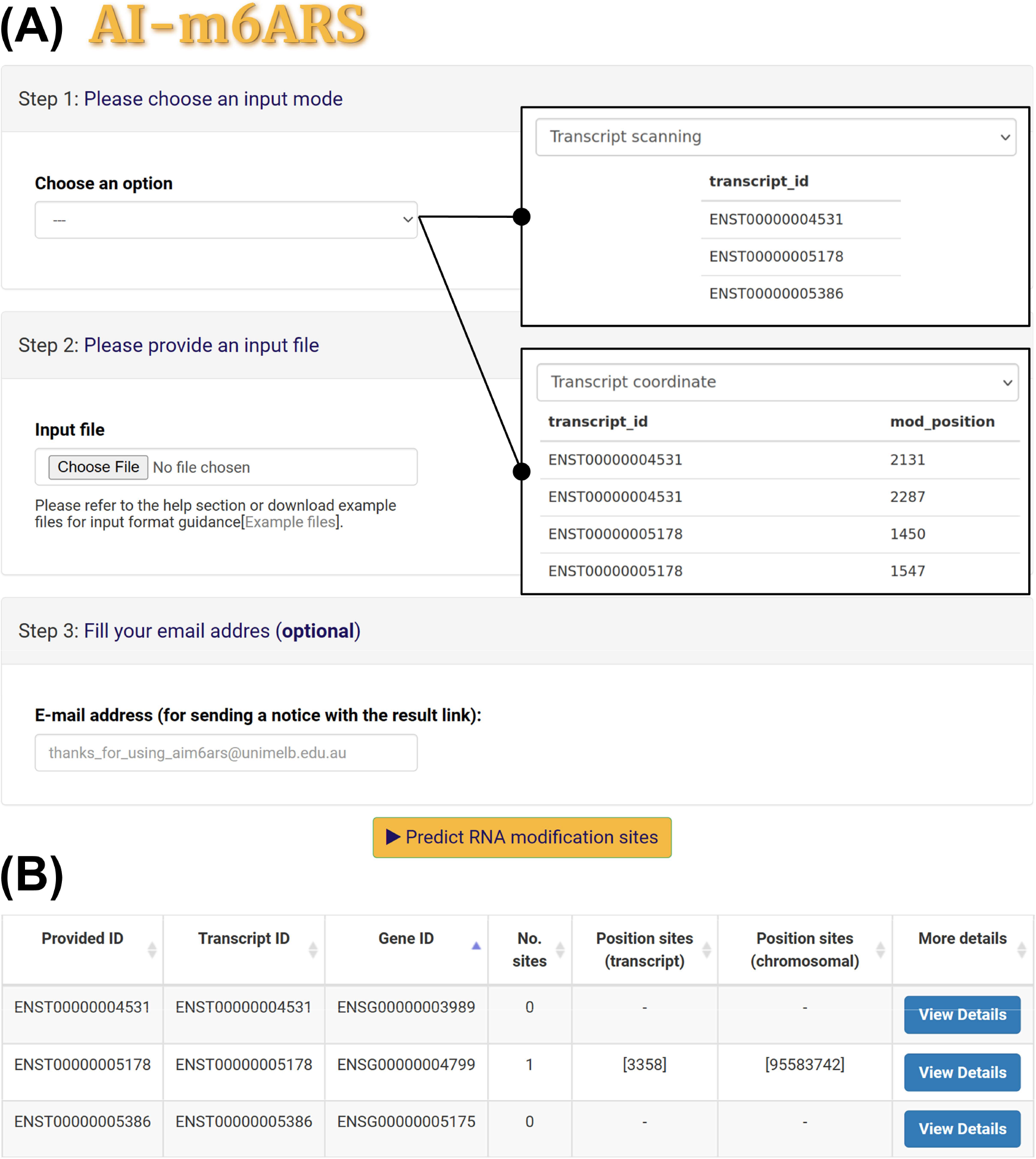
AI-m6ARS web server interface. (A) The AI-m6ARS web server offers two input modes: Transcript scanning and Transcript coordinate. The first mode produces comprehensive predictions for every DRACH motif present in the input transcript. The second mode only generates predictions using input transcripts and provides nucleotide positions. (B) The result of AI-m6ARS is presented in tabular format. The provided information includes the number of predicted m6A sites and their corresponding locations. Additional details, such as the general details of transcripts, predicted probability, DRACH motif sequence, and the distribution of m6A modification sites, can be found in the “View details” section.

## DISCUSSION

m6A methylation plays a crucial role in post-transcriptional gene regulation, exerting major impacts on diverse cellular functions. Multiple studies have established a link between its dysregulation and a diverse array of diseases, such as cancer and neurological disorders (4,14-17,84). The significance of m6A in the progression of diseases not only demonstrates its vital functions but also presents opportunities for developing therapeutic and diagnostic tools (85). Computational methods have been developed to comprehend the intricate landscape of m6A methylation by predicting human m6A methylation sites. These tools address the limitations of traditional experimental methods that struggle to capture all m6A sites in various cellular and genomic contexts, such as different cell types and tissues.

### Strengths

This study presents AI-m6ARS, a novel machine learning model developed for the precise identification of human m6A methylation sites. To address the constraints found in existing computational methods, we have incorporated a wide range of approaches to improve the characterisation of sequence, conservation, and geographical characteristics associated with functional m6A sites, leveraging miCLIP data (27). AI-m6ARS includes an extensive array of sequence features, integrating both one-hot encoding and 25 physicochemical sequence encoding methods provided by iLearn (49,50). To expand the range of m6A site characterisation beyond sequence contexts, a comprehensive literature review was conducted, followed by the integration of updated conservation scores and geographical features. We utilised data preprocessing methods and CD-HIT (55) to improve the quality of our training set. A robust feature selection pipeline was performed to identify and retain the most relevant features. LightGBM, a highly efficient gradient boosting algorithm, was thereafter employed to train a machine learning model (56).

The performance of AI-m6ARs was evaluated through cross-validations and an independent blind test, demonstrating its proficiency in predicting human m6A sites with an AUC, bACC, F1, and MCC of up to 0.86, 0.75, 0.45, and 0.40, respectively (Table 1). When applied to a mouse dataset, AI-m6ARS successfully predicted mouse m6A sites. This reflects the robustness of our pipeline across different species (Table 3). Subsequent analysis indicates that the integration of both human and mouse data significantly improves predictive accuracy in both species, highlighting the advantages of leveraging multi-species data to overcome data limitations (Table 3).

An In-depth model interpretation illustrates the advantages of incorporating sets of features that represent sequence, conservation, and geographical characteristics. Sequence and geographical features are key factors in the model decision-making process. Conservation scores, despite their relatively smaller contribution, still significantly improve model prediction capability (Figure 3 and Table 4). AI-m6ARS demonstrates a slightly improved overall performance when compared to a state-of-the-art model (Table 2). The advantages of our model are the model interpretability of LightGBM algorithms and the availability of a user-friendly web server. The AI-m6ARS web server can be accessed at https://biosig.lab.uq.edu.au/aim6ars/. AI-m6ARS provides accurate predictions for DRACH motifs present in input transcripts, offering valuable insights into the distribution of m6A modification sites (Figure 4 and Figure S3). The capability to prioritise potential m6A sites enable researchers to optimise their experimental efforts, resulting in more efficient and cost-effective validations.

Future directions aim to broaden the scope of AI-m6ARS’s applicability beyond the genomes of humans and mice, while also investigating other essential methylation types. The m6A methylation, abundantly found in eukaryotic mRNA, has been observed in various organisms, such as yeasts, plants, and animals. Each of them exhibits a distinct pattern and unique consensus motif (86). Moreover, a recent finding has reported the existence of m6A methylated sites in bacterial organisms (87). Understanding the evolutionary conservation and divergence of methylation mechanisms and sequence preferences can be facilitated by investigating methylation dynamics across a wide range of species. Additionally, it has the potential to uncover new roles of m6A methylation in different processes.

In addition to m6A, other RNA modifications, such as N1-methyladenosine (m1A) or Inosine (I) (88), also play a vital role in biological processes and have been associated with disease progressions (89). The investigation into the AI-m6ARS pipeline applications in other types of RNA modifications demonstrates significant promise. Highlighting the model’s capability to handle various modification types would underline its adaptability. Conducting a comprehensive literature review is necessary in order to identify the distinct characteristics, such as specific motifs and site deposition mechanisms, associated with each type of modification. The AI-m6ARS pipeline aims to develop multiple models for multiple-species and multiple-methylation data. These models aim to provide a comprehensive understanding of RNA modification, which can be used to develop therapeutic interventions and diagnostic applications for various diseases.

### Feature engineering

This study has demonstrated the significance of employing effective representations of m6A methylation sites to improve the model performance. AI-m6ARS utilises a combination of iFeatures, conservation scores, and geographical features to achieve a good performance. Further improvements can be achieved by employing more effective sequence-context representation or incorporation of supplementary data types, such as the degree of conservation and geographical information of m6A sites.

In a preliminary analysis, we employed RNABERT to encode the sequence pattern of a 41-nucleotide segment containing an m6A site. RNABERT is an RNA-based embedding method that adapts the pre-trained BERT algorithm to non-coding RNA, demonstrating successful applications in structural alignment and clustering (90). Utilising only a set of 4,706 RNABERT features, the model successfully predicts m6A sites from RMBase v2.0 (91) with an AUC of 0.762. Nevertheless, the computationally demanding nature of RNABERT posesd limitations. While efficient for conducting a single prediction or model training, it is not suitable for conducting genome-wide scanning or implementing it within a web server. Furthermore, it is not feasible to pre-generate and store feature values for all possible combinations of 41-nucleotide regions (4^41^ combinations). In the future, we expected that leveraging large language models (LLMs) designed for RNA sequences in a computationally efficient manner could offer a practical solution for more accurate characterisation of functional m6A sites.

### Limitations

One limitation of this study is the quality of the miCLIP dataset. The machine learning models recognise underlying patterns of m6A deposition from experimental datasets to generate accurate predictions. However, the low quality of the dataset may fail to provide a true ground truth to the model. While miCLIP serves as a primary approach for investigating m6A modifications and developing machine learning models, its shortcomings in sensitivity and specificity can limit the model performance and evaluation. It poses a challenge for acquiring true non-m6A sites or negative samples. The lack of m6A at particular DRACH motifs can be ascribed to low sensitivity, making it difficult to distinguish between accurate true negative samples or experimental errors. It is essential to tackle these challenges to improve the reliability and robustness of machine learning models.

DRS, or direct nanopore RNA sequencing, has become a promising alternative technique for identifying m6A sites due to its superior sensitivity and specificity [citation]. DRS offers simultaneous stoichiometric and positional information for multiple modifications. Nevertheless, the determination of m6A sites from DRS raw signals presents difficulties due to subtle variations in signals associated with methylated sites, which are modulated by the sequence context and the variability in translocation rate and pore-to-pore characteristics (92). Several computational models, including xPore (93), Nanocompore (94), and CHEUI (92), have been developed to identify m6A from DRS raw signals with good accuracy.

In future work, we plan to extend our method to handle DRS data. Accordingly, as large amount of DRS data will likely be expanded across various human cell lines and more accurate bioinformatic pipelines, the extensive datasets of reliable m6A and non-m6A sites in different cellular and genomic contexts will be accessible and therefore will be able to be exploited. We plan to create a machine learning model specifically designed to forecast human m6A sites, considering variations in cell lines, the existence of neighbouring methylation, and distinctions between isoforms, as well as their stoichiometry. The advanced models offer the potential to improve the ability to understand the complex mechanisms of m6A modifications.

## DATA AVAILABILITY

The miCLIP datasets used to train AI-m6ARS have been deposited in the web server at https://biosig.lab.uq.edu.au/aim6ars/data.

## SUPPLEMENTARY DATA

Supplementary Data are available at NAR online.

## AUTHOR CONTRIBUTIONS

Korawich Uthayopas: Conceptualization, Data curation, Methodology, Formal analysis, Validation, Visualization, Software, Writing—original draft. Alex G. C. de Sa: Methodology, Supervision, Software, Writing—review & editing. David B. Ascher: Conceptualization, Methodology, Supervision, Writing—review & editing.

## ACKNOWLEDGEMENTS

We wish to thank Azadeh Alavi (RMIT) for discussions about machine learning strategies.

## FUNDING

K.U. was supported by the UQ research scholarship. This research was funded by Investigator Grant from the National Health and Medical Research Council (NHMRC) of Australia [GNT1174405 to D.B.A.] and Victorian Government’s Operational Infrastructure Support Program (in part). Funding for open access charge: National Health and Medical Research Council.

## CONFLICT OF INTEREST

The authors have no relevant financial or nonfinancial interests to disclose.

## Notes

### Competing Interest Statement

The authors have declared no competing interest.

