## Supplementary document 1 for "AI-m6ARS: Machine learning-driven m6A RNA methylation site discovery with integrated sequence, conservation, and geographical descriptors"

### MANUSCRIPT TITLE

### SUPPLEMENTARY LITERATURE REVIEW

#### Experimental methods for an identification of m6A methylation

Antibody-based methods leverage the key concepts of the crosslinking and immunoprecipitation (CLIP) protocol to investigate RNA-protein interactions (1-3). Initially, an anti-m6A antibody is incubated with the sample, followed by UV-crosslinking and protein purification. Proteinase K subsequently breaks down the antibody, resulting in the presence of an amino acid fragment on the RNA base. This can lead to the occurrence of errors during reverse transcription. Computational analysis of the cDNA sequences then identifies the modification site. Two variations of CLIP, miCLIP (2) and $m^6$A-CLIP (3), have been proposed. Enzyme-based techniques, such as MAZTER-seq (4) and m6A-REF-seq (5), exploit the unique properties of the MazF endoribonuclease. The enzyme preferentially cleaves the ACA sequence motif over m6A-CA sites, allowing for the detection of m6A modifications in DRACH motifs through a reduction in MazF cleavage efficiency (4-6).

The DART-seq method (5), based on fusion domains, utilises the fusion of the cytidine deaminase APOBEC1 and m6A-binding YTH domain. This fusion induces deamination near m6A sites, detectable through RNA-seq. This approach successfully identifies m6A sites with a low input requirement and simplified library preparation. On the other hand, DRS has emerged as a highly promising alternative technique for identifying m6A sites through the measurement of changes in ionic current (7,8). One notable advantage of DRS is its capability to provide both stoichiometric and positional information on multiple types of modifications simultaneously. However, the computational analysis of raw signals to infer m6A sites still faces challenges. These include issues such as variance in background current, which reduces detection sensitivity, as well as concerns about the quality and bias of data used as the ground truth in machine learning algorithms (9).

### SUPPLEMENTARY MATERIAL AND METHOD

#### Validation and performance metrics

***k-fold Cross-validation (CV).*** k-fold Cross-validation is a widely used approach for unbiased evaluation of a machine learning model performance. This method ensures that the model is evaluated using an independent dataset that does not overlap with the training dataset. The process starts by partitioning a dataset into k subsets for each fold and then validating using the remaining subsets. In k-fold cross-validation, the performance metric is calculated as the mean of the results obtained in each fold (10,11).

***Area Under the receiver operating characteristic Curve (AUC).*** The Area Under the Curve (AUC) is a common evaluation metric for assessing the performance of machine learning models in classification tasks across different threshold values. The area under the receiver operating characteristic (ROC) curve, which is plotted with the True Positive Rate (TPR) and False Positive Rate (FPR), represents the model's capability to differentiate between different classes. AUC values range from 0 to 1, with an exceptional model achieving AUC values close to 1. An AUC of 0.5 indicates that the model lacks the ability to differentiate between different classes (10,11).

***Balanced accuracy (bACC).*** The bACC metric is an additional measure used to assess the classification model. The calculation involves determining the mean value of the True Positive Rate (TPR), also known as Sensitivity, and the True Negative Rate (TNR), also known as Specificity (10,11).

***Matthews Correlation Coefficient (MCC).*** MCC is a statistical metric employed for the evaluation of binary classification models by assessing the degree of correlation between predicted and actual values (10,11). MCC values range from -1 to 1, where a perfect model is assigned a score of 1. The calculation of MCC is performed using the following formula:

$$\boldsymbol{MCC} = \frac{\boldsymbol{TP} \times \boldsymbol{TN} - \boldsymbol{FP} \times \boldsymbol{FN}}{\sqrt{}(\boldsymbol{TP} + \boldsymbol{FP})(\boldsymbol{TP} + \boldsymbol{FN})(\boldsymbol{TN} + \boldsymbol{FP})(\boldsymbol{TN} + \boldsymbol{FN})}$$

Where TP, TN, FP, and FN refer to True positive, True negative, False positive, and False negative.

***Precision.*** Precision is a frequently used metric for quantifying the performance of a model. It refers to the proportion of true positive sample predictions that were accurately predicted by the model (10,11).

***Recall, Sensitivity, or True Positive Rate (TPR).*** The term "recall" or "sensitivity" refers to the capability of a model to accurately identify positive samples. The metric is determined by dividing the total count of true positives by the total count of predicted positives (10,11).

***Specificity or True Negative Rate (TNR).*** The concept of specificity relates to the capability of a model to accurately classify negative samples as negative samples. The metric can be calculated by dividing the total count of true negatives by the total count of predicted negatives (10,11).

***F1-score.*** The F1-score is a statistical measure that integrates the precision and recall of a classifier by employing the harmonic mean. Its main purpose is to assess the effectiveness of binary classification. The achievement of a high F1-score is dependent on the classifier's predictive power to achieve both high precision and high recall (10,11).

**Feature preprocessing and data scaling**

Feature preprocessing plays a crucial role in the machine learning pipeline as it converts raw data into an appropriate format for training accurate machine learning models [10]. The essential steps involve handling missing data, addressing outliers, scaling feature values, and encoding categorical variables. Scikit-learn, a standard Python library for machine learning, provides six different types of scalers in the preprocessing module (12). Each scaler is specifically designed to address unique challenges in feature preprocessing. The choice of a suitable scaler relies on the distinctive characteristics of the dataset and the requirements of machine learning algorithms.

***StandardScaler.*** The technique referred to as the StandardScaler, or Z-score normalisation, is employed to scale features in order to achieve a mean of 0 and a standard deviation (SD) of 1. Although this approach demonstrates utility in achieving a standardised distribution, it is unable to guarantee balanced feature scales in the presence of outliers (10).

***MinMaxScaler.*** The MinMaxScaler method is utilised to rescale feature values within the range of 0 to 1. In this process, the minimum value is subtracted from each feature, followed by division by the range, which is defined as the difference between the maximum and minimum values. The MinMaxScaler algorithm shows susceptibility to the presence of outliers (10).

***MaxAbsScaler.*** The MaxAbsScaler is a method used to transform feature values into a range spanning from -1 to 1. The procedure consists of the division of each feature by its maximum absolute value. Although MaxAbsScaler exhibits lower sensitivity to outliers in comparison to certain alternative methods, it remains susceptible to the impact of large outliers.

***PowerTranformer.*** This method, known as PowerTransformer, employs a power transformation on each feature in order to produce a Gaussian distribution for the data. This transformation serves to stabilise variance and reduce skewness (10).

***RobustScaler.*** The RobustScaler utilises percentiles to perform feature scaling, making it well-suited for effectively dealing with large outliers. The scaling method described is particularly suitable for datasets that include outliers, as it provides a high level of robustness even when extreme values are present (10).

***QuantileTranformer.*** In order to align the probability density function of each feature with either a uniform or normal distribution, the QuantileTransformer employs a nonlinear transformation. This approach holds significant advantages in situations where machine learning algorithms require normality. Furthermore, it effectively addresses the effects of outliers and skewed distributions within the dataset (10).

**Feature selection**

***Variance Threshold.*** The variance threshold algorithm is a feature selection method aimed to eliminate features that exhibit low variance (12,13). Features with low variance indicate a limited amount of variation within the dataset, making them less informative for training machine learning models. The procedure involves the removal of low-variance features through the establishment of a predefined threshold. It can be employed using the scikit-learn library in Python (12).

***Correlation Threshold.*** The correlation threshold is a technique designed to eliminate features that demonstrate a significantly strong correlation with one another (13). The correlation threshold employs a predetermined threshold for the correlation coefficient between pairs of features. To mitigate redundancy in the dataset, it is possible to eliminate one of the features when the correlation between them surpasses a certain threshold. The scikit-learn library in Python can be utilised for correlation thresholds (12).

***Boruta.*** Boruta is an algorithm for feature selection that has been specifically developed to identify and choose relevant features from a given set of features (14). The Random Forest classifier is utilised by Boruta as its base model. Boruta generates shadow features for each feature in the dataset by rearranging the values of that specific feature. The shadow features function as a benchmark for evaluating the impact of the original features on the model. An original feature is considered important if its feature importance statistically surpasses that of its shadow feature. The procedure is iterated multiple times, during which unnecessary features are eliminated. One benefit of Boruta is its independence from a particular data distribution and its simplicity. The BorutaPy package in Python is used for the implementation of Boruta (14).

**Machine learning algorithms**

***Light Gradient Boosting Machine (LightGBM).*** LightGBM is a gradient boosting framework developed by Microsoft to facilitate the efficient training of large datasets (15). It employs a tree-based learning algorithm and leaf-wise growth strategy for node splitting, decreasing memory usage, and enhancing training efficiency. This characteristic makes it highly effective in managing large datasets.

***Gradient Boosting (GB).*** Gradient boosting is a machine learning method that constructs a predictive model through the integration of numerous weak learners, typically composed of decision trees (16). The model iteratively corrects errors made by its preceding iterations, thereby iteratively enhancing the accuracy of its predictions.

***Explainable Boosting Machine (EBM).*** EBM stands out as an interpretable machine learning model designed to provide insights into its decision-making process. By combining the interpretability of generalised additive models with boosting algorithms, EBM offers a balance between model complexity and transparency (17).

**Extreme Gradient Boosting (XGBoost).** XGBoost is a highly efficient and scalable implementation of gradient boosting (18). It incorporates regularisation into its objective function to prevent overfitting. XGBoost also allows a parallel implementation to accelerate training. The versatility of XGBoost and its high-performance capabilities have made it a popular choice in machine learning competitions.

***Adaptive Boosting (ADABoost).*** The ADABoost algorithm constructs a robust model through a combination of numerous weak learners (19). The algorithm assigns varying weights to instances, taking into account their classification performance in prior models, with a particular emphasis on instances that were previously misclassified. ADABoost is highly effective for binary classification problems due to its ability to dynamically adapt to the complexity of the data.

***Random Forest (RF).*** The Random Forest algorithm is an ensemble learning technique that leverages the construction of numerous decision trees during the training process (20). It enhances predictive performance and mitigates overfitting by incorporating randomness through the selection of a subset of features for each tree. Random forests are highly adaptable and extensively employed across various domains.

***Extremely Randomised Trees (Extra Trees).*** Extra Trees is another ensemble learning algorithm similar to Random Forest. The algorithm constructs multiple decision trees with random splits and subsequently selects the trees that exhibit the best performance to form the ensemble (21). The addition of randomness in a feature selection process during tree construction improves the model's robustness, contributing to the favourable performance achieved by Extra Trees.

***Multilayer perceptron (MLP).*** MLP is a type of artificial neural network characterised by multiple layers of interconnected nodes, known as neurons (22). This architecture allows MLPs to recognise complex patterns in data. Although MLPs offer excellent predictive capabilities, they may require cautious hyperparameter tuning. MLP is commonly used in various complex tasks, including image recognition, natural language processing, and other applications.

***K-Nearest Neighbors (KNN).*** KNN is a straightforward machine learning algorithm that predicts the results by considering the majority class or the average of the k-nearest data points in the feature space [23]. The simplicity of KNN makes it easy to understand and implement. Nevertheless, the performance of the system can be influenced by the selection of the distance metric and the number of neighbours.

***LogisticRegression.*** Logistic regression is a linear model employed for binary classification. (23) This method employs the logistic function to estimate the likelihood that a given data point is associated with a specific class. Frequently employed as a baseline model in classification tasks.

**TABLE AND FIGURES LEGENDS**

**Table S1.** Overview of computational methods for predicting human m6A methylation sites.

| Models | Publication date | Features | Algorithm | Citation | Availability | URLs | Accessibility |
|---|---|---|---|---|---|---|---|
| MethyRNA | Jan, 2016 | • Physico-chemical-based one-hot encoding[a]<br>• Cumulative nucleotide frequency | Support Vector Machine (SVM) | (24) | Web server | https://lin.uestc.edu.cn/server/methyrna | No<br>(As of 25 Feb, 2024) |
| SRAMP<br>(Sequence-based RNA Adenosine Methylation Site Predictor) | Jun, 2016 | • One-hot encoding<br>• K-nearest neighbour (KNN)<br>• Nucleotide pair spectrum<br>• RNAfold predicted secondary structure | Random Forest (RF) | (25) | Web server, Source code | https://www.cuilab.cn/sramp/ | Yes |

| | | | | | | | |
|---|---|---|---|---|---|---|---|
| RNAMethPre | Oct, 2016 | • One-hot encoding<br>• K-mer composition<br>• Relative site position in mRNA<br>• RNAfold Minimum Free Energy (MFE) | Support Vector Machine (SVM) | (26) | Web server | https://bioinfo.tsinghua.edu.cn/RNAMethPre/index.html | No (As of 25 Feb, 2024) |
| RAM-NPPS (RNA $N^6$-Adenosine Methylation - Nucleotide Pair Position Specificity) | Apr, 2017 | • Nucleotide Pair Position Specificity (NPPS) | Support Vector Machine (SVM) | (27) | Web server | http://server.malab.cn/RAM-NPPS/ | No (As of 25 Feb, 2024) |
| HMPre | Aug, 2018 | • One-hot encoding<br>• Chemical Property with Density (CPD) | eXtreme Gradient Boosting | (28) | Source code | https://github.com/Zhixun-Zhao/HMpre | Yes |

| | | | | | | | |
|---|---|---|---|---|---|---|---|
| | | • K-mer composition<br>• Site-location features<br>• Entropy features<br>• SNP features | (XGBoost) | | | | |
| BERMP<br>(BGRU-base Ensemble RNA Methylation site Predictor) | Sep, 2018 | • Enhanced Nucleic Acid Composition (ENAC) | Bidirectional Gated Recurrent Unit Neural Network (BGRU) | (29) | Source code | http://www.bioinfogo.org/bermp | No<br>(As of 25 Feb, 2024) |
| Gene2vec<br>(Gene subsequence to embedding | Feb, 2019 | • One-hot encoding<br>• Neighbouring methylation state encoding<br>• RNA word embedding | Convolution neural networks (CNNs) | (30) | Web server | https://server.malab.cn/Gene2vec/ | No<br>(As of 25 Feb, 2024) |

| | | | | | | | |
|---|---|---|---|---|---|---|---|
| vector) | | • Gene2vec | | | | | |
| WHISTLE<br><br>(Whole-transcriptome $m^6A$ site prediction from multiple genomic features) | Apr, 2019 | • Physico-chemical-based one-hot encoding[a]<br>• Cumulative nucleotide frequency<br>• 35 Genome-based features[b] | Support vector machine (SVM) | (31) | Prediction results | http://180.208.58.19/whistle/index.html | Yes |
| DeepM6ASeq | Dec, 2019 | • One-hot encoding | Convolution neural networks (CNNs) | (32) | Source code | https://github.com/rreybeyb/DeepM6ASeq | Yes |
| WITMSG<br><br>(Whole-intronome $m^6A$ methylation sites | Jan, 2020 | • Physico-chemical-based one-hot encoding[a]<br>• Cumulative nucleotide frequency | Random Forest (RF) | (33) | Prediction results | http://rnamd.com/intron/ | Yes |

| | | | | | | | |
|---|---|---|---|---|---|---|---|
| prediction by combining sequence features with genomic features) | | • 60 Genome-based features[b] | | | | | |
| LITHOPHONE<br><br>(Long noncoding RNA methylation sites prediction from sequence characteristics and genomic information with an ensemble predictor) | Jun, 2020 | • Physico-chemical-based one-hot encoding[a]<br>• Cumulative nucleotide frequency<br>• 60 Genome-based features[b] | Random Forest (RF) | (34) | Prediction results | http://180.208.58.19/lith/ | No<br>(As of 25 Feb, 2024) |
| DeepPromise<br><br>(Deep CNN-based | Sep, 2020 | • One-hot encoding | Convolutional neural | (35) | Web server | https://deeppromise.erc.monas | No<br>(As of 1 Apr, |

| | | | | | | | |
|---|---|---|---|---|---|---|---|
| Predictor of RNA modification sites) | | • RNA word embedding<br>• Enhanced nucleic acid composition (ENAC) | network (CNN) | | | h.edu/ | 2024) |
| EDLm$^6$APred (Ensemble deep learning m$^6$A site predictor) | May, 2021 | • One-hot encoding<br>• RNA word embedding<br>• Word2vec | Bidirectional LSTM (BiLSTM) | (36) | Web server | https://www.xjtlu.edu.cn/biologicalsciences/EDLm6APred | No (As of 25 Feb, 2024) |
| MultiRM (Attention-based Multi-label neural network approach for integrated prediction and interpretation of RNA modifications | Jun, 2021 | • One-hot encoding<br>• Word2vec<br>• Hidden Markov Model (HMM) | Long short-term memory (LSTM) | (37) | Web server<br><br>Source code | www.xjtlu.edu.cn/biologicalsciences/multirm<br><br>https://github.com/Tsedao/MultiRM | No (As of 25 Feb, 2024)<br><br>Yes |

| | | | | | | | |
|---|---|---|---|---|---|---|---|
| from the primary RNA sequence) | | | | | | | |
| TS-m6A-DL<br><br>(Tissue-specific identification of m6A sites using a universal deep learning model) | Aug, 2021 | ● One-hot encoding | Convolutional neural network (CNN) | (38) | Web server | nsclbio.jbnu.ac.kr/tools/TS-m6A-DL/ | Yes |
| HSM6AP<br><br>(High-precision predictor for the Homo sapiens $m^6A$ based on multiple weights and feature stitching) | Nov, 2021 | ● 4 Sequence-based features - K-mer composition, Composition of *k*-spaced nucleic acid pairs (CKSNAP), Di-Nucleotide Composition (DNC), Mismatch<br><br>● 5 Physico-chemical features - Nucleotide Chemical Property (NCP), Parallel Correlation Pseudo Dinucleotide Composition | Extreme gradient boosting (XGBoost) | (39) | Web server<br><br>Source code | http://120.27.24.199:10501/HSM6AP/WebServer<br><br>https://github.c | No<br><br>(As of 3 Apr, 2024) |

| | | | | | | | |
|---|---|---|---|---|---|---|---|
| | | (PCPseDNC), Parallel Correlation Pseudo Trinucleotide Composition (PCPseTNC), Series Correlation Pseudo Dinucleotide Composition (SCPseDNC), Series Correlation Pseudo Trinucleotide Composition (SCPseTNC)<br>• 64 Genome-based features[b] | | | | om/lijingtju/HSm6AP | No<br>(As of 3 Apr, 2024) |
| GepSe<br>(Geography plus Sequences) | Sep, 2022 | • One-hot encoding<br>• Geographic representation of transcript as vectors (Geo2Vec) - landmarkTX, gridTX, and chunkTX | Convolutional neural network (CNN) | (40) | Web Server<br><br>Source code | https://www.xjtlu.edu.cn/biologicalsciences/geo2vec<br><br>https://github.com/daiyun02211/Geo2vec | No<br><br>Yes |

| | | | | | | | |
|---|---|---|---|---|---|---|---|
| DL-m6A (Identification of m6A sites in mammals using deep learning based on different encoding schemes) | Apr, 2023 | • One-hot encoding<br>• Nucleotide Chemical Property (NCP)<br>• Nucleotide Density (ND)<br>• Electron-ion interaction potential (EIIP) | Convolutional neural network (CNN) | (41) | Web server | nsclbio.jbnu.ac.kr/tools/DL-m6A/ | Yes |
| GR-m6A | Sep, 2023 | • SMILES strings | Multilayer perceptron (MLP) | (42) | Source code | https://github.com/YingLiangjxau/GR-m6A | Yes |

[a]Physico-chemical-based one-hot encoding categorises four types of nucleotides using three features: a ring number, a strength of hydrogen bonds, and functional groups. [b]Genome-based features are designed to encompass various types of genomic information, such as positional data in transcripts, conservation scores, RNA secondary structures, and distances to splicing junction.

**Table S2.** 6,396 features used to model human m6A methylation sites in AI-m6ARS.

| Category | Subcategory | Description | Number of features | Sources / Tools |
|---|---|---|---|---|
| One-hot encoding | - | The process of one-hot encoding involves the transformation of four distinct types of nucleobases into binary vectors. This process produces a total of 164 unique features, which correspond to 41-nucleotide sequences that are centred around m6A methylation sites. | 164 | Python script |
| iFeatures | - | iFeatures comprise a set of sequence-derived features acquired from the iLearn Python toolkit. These features are frequently employed to construct a diverse array of features, effectively encoding the physicochemical characteristics present in DNA, RNA, or protein sequences.<br><br>A total of 25 different RNA sequence encoding schemes are utilised. They include: 1) Kmer type 1 2) Mismatch 3) Subsequence 4) ENAC 5) ANF 6) Binary 7) PS2 8) PS3 9) CKSNAP type 1 10) NCP 11) ASDC 12) DBE 13) LPDF 14) DPCP type 2 15) Z curve 9 bit 16) Z curve 12 bit 17) Z curve 36 bit 18) NMBroto 19) Moran 20) Geary 21) DACC 22) DCC 23) PseKNC 24) PC-PseDNC 25) SC-PseDNC | 4,591 | iLearn library (43) |

| | | | | |
|---|---|---|---|---|
| | | For detailed descriptions of feature descriptors for nucleotide sequences, please refers to online manual in iLearnPlus website (https://ilearnplus.erc.monash.edu/docs/iLearnPlus_manual.pdf) | | |
| Conservation scores | 100-way PhastCons | In order to characterise m6A sites in terms of evolutionary constraints, conservation scores are utilised to quantify the level of conservation within the m6A methylation site, DRACH motif, and adjacent region.<br><br>The PhastCons algorithm is a computational method based on a hidden Markov model. Its purpose is to estimate the probability of a particular nucleotide being associated with a conserved element.<br><br>The UCSC Genome Browser was used to obtain 100-way PhastCons scores from a multiple sequence alignment of 100 vertebrate species. A total of nine features were generated, encompassing:<br><br>1) A conservation score of the modification site<br>2) A conservation score of 1st position in DRACH motif (D)<br>3) A conservation score of 2nd position in DRACH motif (R)<br>4) A conservation score of 4th position in DRACH motif (C)<br>5) A conservation score of 5th position in DRACH motif (H) | 9 | UCSC Genome Browser (44) |

| | | | | |
|---|---|---|---|---|
| | | 6) A mean value of conservation scores in DRACH motifs<br>7) A mean value of conservation scores from 5-nt upstream to 5-nt downstream of modification site.<br>8) A mean value of conservation scores from 10-nt upstream to 10-nt downstream of modification site.<br>9) A mean value of conservation scores from 20-nt upstream to 20-nt | | |
| | 30-way PhastCons | In order to characterise m6A sites in terms of evolutionary constraints, conservation scores are utilised to quantify the level of conservation within the m6A methylation site, DRACH motif, and adjacent region.<br>The PhastCons algorithm is a computational method based on a hidden Markov model. Its purpose is to estimate the probability of a particular nucleotide being associated with a conserved element.<br>The UCSC Genome Browser was used to obtain 30-way PhastCons scores from a multiple sequence alignment of 30 vertebrate species. A total of nine features were generated, encompassing:<br>1) A conservation score of the modification site<br>2) A conservation score of 1$^{st}$ position in DRACH motif (D)<br>3) A conservation score of 2$^{nd}$ position in DRACH motif (R)<br>4) A conservation score of 4$^{th}$ position in DRACH motif (C)<br>5) A conservation score of 5$^{th}$ position in DRACH motif (H) | 9 | UCSC Genome Browser (44) |

| | | | | |
|---|---|---|---|---|
| | | 6) A mean value of conservation scores in DRACH motifs<br>7) A mean value of conservation scores from 5-nt upstream to 5-nt downstream of modification site.<br>8) A mean value of conservation scores from 10-nt upstream to 10-nt downstream of modification site.<br>9) A mean value of conservation scores from 20-nt upstream to 20-nt downstream of modification site. | | |
| | 100-way PhyloP | Conservation scores are employed to quantify the degree of conservation within the m6A methylation site, DRACH motif, and surrounding region, with the aim of characterising m6A sites in terms of evolutionary constraints.<br>PhyloP assesses the evolutionary conservation of particular nucleotide positions through a comparative analysis with other species. It calculates p-values based on a sequence alignment and a mode of neutral evolution using independent hypothesis tests.<br>Positive scores indicate conservation at the site, while negative scores indicate rapid evolution of the site.<br>The 100-way PhyloP scores were acquired from a multiple sequence alignment of 100 vertebrate species retrieved from the UCSC Genome Browser. Nine features were generated in total, comprising:<br>1) A conservation score of the modification site<br>2) A conservation score of $1^{st}$ position in DRACH motif (D) | 9 | UCSC Genome Browser (44) |

| | | | | |
|---|---|---|---|---|
| | | 3) A conservation score of $2^{nd}$ position in DRACH motif (R)<br>4) A conservation score of $4^{th}$ position in DRACH motif (C)<br>5) A conservation score of $5^{th}$ position in DRACH motif (H)<br>6) A mean value of conservation scores in DRACH motifs<br>7) A mean value of conservation scores from 5-nt upstream to 5-nt downstream of modification site.<br>8) A mean value of conservation scores from 10-nt upstream to 10-nt downstream of modification site.<br>9) A mean value of conservation scores from 20-nt upstream to 20-nt downstream of modification site. | | |
| | 30-way PhyloP | Conservation scores are employed to quantify the degree of conservation within the m6A methylation site, DRACH motif, and surrounding region, with the aim of characterising m6A sites in terms of evolutionary constraints.<br><br>PhyloP assesses the evolutionary conservation of particular nucleotide positions through a comparative analysis with other species. It calculates p-values based on a sequence alignment and a mode of neutral evolution using independent hypothesis tests.<br><br>Positive scores indicate conservation at the site, while negative scores indicate rapid evolution of the site.<br><br>The 30-way PhyloP scores were acquired from a multiple sequence alignment of 30 vertebrate species retrieved from the UCSC Genome Browser. Nine features were | 9 | UCSC Genome Browser (44) |

| | | | | |
|---|---|---|---|---|
| | | generated in total, comprising:<br>1) A conservation score of the modification site<br>2) A conservation score of 1st position in DRACH motif (D)<br>3) A conservation score of 2nd position in DRACH motif (R)<br>4) A conservation score of 4th position in DRACH motif (C)<br>5) A conservation score of 5th position in DRACH motif (H)<br>6) A mean value of conservation scores in DRACH motifs<br>7) A mean value of conservation scores from 5-nt upstream to 5-nt downstream of modification site.<br>8) A mean value of conservation scores from 10-nt upstream to 10-nt downstream of modification site.<br>9) A mean value of conservation scores from 20-nt upstream to 20-nt downstream of modification site. | | |
| Geographical features | Distance-based | The primary objective of geographical features is to utilise positional information associated with m6A methylation sites in order to improve the characterisation of m6A site deposition.<br>Positional information of m6A sites within transcripts is captured by Distance features, which encompass the distances to geographical landmarks such as the start and end of the transcript, exons, splice sites, 5′-UTR, and 3′-UTR.<br>A set of 15 features derived from distance-based geographical features are created, | 15 | Python script |

| | | |
|---|---|---|
| | | consisting of:<br>1) Transcript length<br>2) Distance to the start of a transcript (always positive)<br>3) Distance to the end of a transcript (always positive)<br>4) Distance to the end of 5′-UTR (always positive)<br>5) Distance to the start of 3′-UTR (always positive)<br>6) Distance to the start of current exon (positive if a site resides in exon)<br>7) Distance to the end of current exon (positive if a site resides in exon)<br>8) Distance to the closest splice site (positive if a splice site exists)<br>9) Is transcript having 5′-UTR<br>10) Is transcript having 3′-UTR<br>11) Is modification site in exon (Y/N)<br>12) Is modification site in 5′-UTR (Y/N)<br>13) Is modification site in CDS (Y/N)<br>14) Is modification site in Intron (Y/N)<br>15) Is modification site in 3′-UTR (Y/N) |

| | | | | |
|---|---|---|---|---|
| | Fragment-based | The primary objective of geographical features is to utilise positional information associated with m6A methylation sites in order to improve the characterisation of m6A site deposition.<br><br>The transcript was partitioned into a specific number of fragments, with each fragment being denoted by nine distinct features, as outlined in the following list.<br><br>1) Number of the fragment<br>2) Length of fragment<br>3) Index of Fragment<br>4) Is m6A resides within this fragment<br>5) Exon composition (0 to 1)<br>6) 5'-UTR composition (0 to 1)<br>7) CDS composition (0 to 1)<br>8) Intron composition (0 to 1)<br>9) 3'-UTR composition (0 to 1)<br><br>These features indicate the composition of particular regions within the fragment, determined by dividing the number of nucleotides in those regions by the length of the fragment. The regions of interest include the exon, 5'-UTR, coding sequences (CDS), introns, and 3'-UTR. The alteration of fragment numbers (8, 16, 32, and 64) enables the generation of features at varying levels of detail, thereby enabling the model to acquire knowledge about the specific context of each fragment and the relative location of the | 1080 | Python script |

| | | | | |
|---|---|---|---|---|
| | | target within the transcripts. | | |
| | Region-based | The primary objective of geographical features is to utilise positional information associated with m6A methylation sites in order to improve the characterisation of m6A site deposition.<br><br>At the regional level, region features are utilised to capture various characteristics, including width, composition, and relative order. The transcripts were divided into a number of regions according to their region type, including exon, 5′-UTR, CDS, intron, and 3′-UTR. The presence of the m6A site, region length, and region type are encoded by 10 features for each region, as shown in the following list.<br><br>1) The region index (-25 to 25)<br>2) Length of a region (nts)<br>3) A proportion of a region length to whole sequence (nts/nts)<br>4) -log2 value of a proportion of a region length to whole sequence (nts/nts)<br>5) A presence of the m6A site within the region. (Y/N)<br>6) Is a region 5′-UTR<br>7) Is a region CDS<br>8) Is a region Intron<br>9) Is a region 3′-UTR<br>10) Is a region exon<br><br>In consideration of the diverse number of distinct regions observed in transcripts, the | 510 | Python script |

| | | |
|---|---|---|
| | | technique of zero-padding was utilised for simpler transcripts, while trimming was employed for more complex transcripts. The m6A site is located at the centre of the feature matrix during feature generation, with a width of 1. |

**Table S4.** 175 selected features implemented in AI-m6ARS.

| No. | Feature name | Feature type | No. | Feature name | Feature type |
|---|---|---|---|---|---|
| 1 | Mismatch_AAA | iFeatures | 89 | Distance-based geo 4 | Geographical features |
| 2 | Mismatch_AAT | iFeatures | 90 | Distance-based geo 5 | Geographical features |
| 3 | Mismatch_ACA | iFeatures | 91 | Distance-based geo 6 | Geographical features |
| 4 | Mismatch_ACG | iFeatures | 92 | Distance-based geo 7 | Geographical features |
| 5 | Mismatch_AGA | iFeatures | 93 | Distance-based geo 8 | Geographical features |
| 6 | Mismatch_AGC | iFeatures | 94 | Fragment-based geo 1073 | Geographical features |
| 7 | Mismatch_AGG | iFeatures | 95 | Region-based geo 42 | Geographical features |
| 8 | Mismatch_AGT | iFeatures | 96 | Region-based geo 62 | Geographical features |
| 9 | Mismatch_ATA | iFeatures | 97 | Region-based geo 64 | Geographical features |
| 10 | Mismatch_ATC | iFeatures | 98 | Region-based geo 72 | Geographical features |
| 11 | Mismatch_ATT | iFeatures | 99 | Region-based geo 74 | Geographical features |
| 12 | Mismatch_CCA | iFeatures | 100 | Region-based geo 82 | Geographical features |
| 13 | Mismatch_CCC | iFeatures | 101 | Region-based geo 84 | Geographical features |

| | | | | | |
|---|---|---|---|---|---|
| 14 | Mismatch_CCG | iFeatures | 102 | Region-based geo 92 | Geographical features |
| 15 | Mismatch_CGA | iFeatures | 103 | Region-based geo 94 | Geographical features |
| 16 | Mismatch_CGC | iFeatures | 104 | Region-based geo 102 | Geographical features |
| 17 | Mismatch_CGG | iFeatures | 105 | Region-based geo 104 | Geographical features |
| 18 | Mismatch_CGT | iFeatures | 106 | Region-based geo 112 | Geographical features |
| 19 | Mismatch_CTT | iFeatures | 107 | Region-based geo 114 | Geographical features |
| 20 | Mismatch_GAA | iFeatures | 108 | Region-based geo 122 | Geographical features |
| 21 | Mismatch_GAC | iFeatures | 109 | Region-based geo 124 | Geographical features |
| 22 | Mismatch_GAG | iFeatures | 110 | Region-based geo 132 | Geographical features |
| 23 | Mismatch_GCA | iFeatures | 111 | Region-based geo 134 | Geographical features |
| 24 | Mismatch_GCC | iFeatures | 112 | Region-based geo 142 | Geographical features |
| 25 | Mismatch_GCG | iFeatures | 113 | Region-based geo 144 | Geographical features |
| 26 | Mismatch_GGA | iFeatures | 114 | Region-based geo 152 | Geographical features |
| 27 | Mismatch_GGC | iFeatures | 115 | Region-based geo 154 | Geographical features |

| | | | | | |
|---|---|---|---|---|---|
| 28 | Mismatch_GGG | iFeatures | 116 | Region-based geo 162 | Geographical features |
| 29 | Mismatch_GGT | iFeatures | 117 | Region-based geo 164 | Geographical features |
| 30 | Mismatch_GTT | iFeatures | 118 | Region-based geo 172 | Geographical features |
| 31 | Mismatch_TAA | iFeatures | 119 | Region-based geo 174 | Geographical features |
| 32 | Mismatch_TAT | iFeatures | 120 | Region-based geo 182 | Geographical features |
| 33 | Mismatch_TCC | iFeatures | 121 | Region-based geo 184 | Geographical features |
| 34 | Mismatch_TGA | iFeatures | 122 | Region-based geo 192 | Geographical features |
| 35 | Mismatch_TGT | iFeatures | 123 | Region-based geo 194 | Geographical features |
| 36 | Mismatch_TTA | iFeatures | 124 | Region-based geo 202 | Geographical features |
| 37 | Mismatch_TTC | iFeatures | 125 | Region-based geo 204 | Geographical features |
| 38 | Mismatch_TTT | iFeatures | 126 | Region-based geo 212 | Geographical features |
| 39 | Subsequence_AAA | iFeatures | 127 | Region-based geo 214 | Geographical features |
| 40 | Subsequence_AGA | iFeatures | 128 | Region-based geo 222 | Geographical features |
| 41 | Subsequence_CCC | iFeatures | 129 | Region-based geo 224 | Geographical features |

| | | | | | |
|---|---|---|---|---|---|
| 42 | Subsequence_GAA | iFeatures | 130 | Region-based geo 232 | Geographical features |
| 43 | CKSNAP_AA.gap0 | iFeatures | 131 | Region-based geo 234 | Geographical features |
| 44 | CKSNAP_AC.gap0 | iFeatures | 132 | Region-based geo 241 | Geographical features |
| 45 | CKSNAP_CC.gap0 | iFeatures | 133 | Region-based geo 242 | Geographical features |
| 46 | CKSNAP_CG.gap0 | iFeatures | 134 | Region-based geo 244 | Geographical features |
| 47 | CKSNAP_CT.gap0 | iFeatures | 135 | Region-based geo 254 | Geographical features |
| 48 | CKSNAP_GA.gap0 | iFeatures | 136 | Region-based geo 262 | Geographical features |
| 49 | CKSNAP_GT.gap0 | iFeatures | 137 | Region-based geo 264 | Geographical features |
| 50 | CKSNAP_TA.gap0 | iFeatures | 138 | Region-based geo 272 | Geographical features |
| 51 | CKSNAP_TT.gap0 | iFeatures | 139 | Region-based geo 274 | Geographical features |
| 52 | CKSNAP_AA.gap1 | iFeatures | 140 | Region-based geo 282 | Geographical features |
| 53 | CKSNAP_AC.gap1 | iFeatures | 141 | Region-based geo 284 | Geographical features |
| 54 | CKSNAP_AT.gap1 | iFeatures | 142 | Region-based geo 292 | Geographical features |
| 55 | CKSNAP_CC.gap1 | iFeatures | 143 | Region-based geo 294 | Geographical features |

| | | | | | |
|---|---|---|---|---|---|
| 56 | CKSNAP_GA.gap1 | iFeatures | 144 | Region-based geo 302 | Geographical features |
| 57 | CKSNAP_GC.gap1 | iFeatures | 145 | Region-based geo 304 | Geographical features |
| 58 | CKSNAP_GT.gap1 | iFeatures | 146 | Region-based geo 312 | Geographical features |
| 59 | CKSNAP_TT.gap1 | iFeatures | 147 | Region-based geo 314 | Geographical features |
| 60 | CKSNAP_AG.gap2 | iFeatures | 148 | Region-based geo 322 | Geographical features |
| 61 | CKSNAP_CC.gap2 | iFeatures | 149 | Region-based geo 324 | Geographical features |
| 62 | CKSNAP_CG.gap2 | iFeatures | 150 | Region-based geo 332 | Geographical features |
| 63 | CKSNAP_GC.gap2 | iFeatures | 151 | Region-based geo 334 | Geographical features |
| 64 | CKSNAP_GG.gap2 | iFeatures | 152 | Region-based geo 342 | Geographical features |
| 65 | CKSNAP_TA.gap2 | iFeatures | 153 | Region-based geo 344 | Geographical features |
| 66 | CKSNAP_TT.gap2 | iFeatures | 154 | Region-based geo 352 | Geographical features |
| 67 | CKSNAP_AA.gap3 | iFeatures | 155 | Region-based geo 354 | Geographical features |
| 68 | CKSNAP_AT.gap3 | iFeatures | 156 | Region-based geo 362 | Geographical features |
| 69 | CKSNAP_CC.gap3 | iFeatures | 157 | Region-based geo 364 | Geographical features |

| | | | | | |
|---|---|---|---|---|---|
| 70 | CKSNAP_GC.gap3 | iFeatures | 158 | Region-based geo 372 | Geographical features |
| 71 | CKSNAP_TT.gap3 | iFeatures | 159 | Region-based geo 374 | Geographical features |
| 72 | DPCP2_Shift (RNA)_pos10 | iFeatures | 160 | Region-based geo 382 | Geographical features |
| 73 | DPCP2_Slide (RNA)_pos10 | iFeatures | 161 | Region-based geo 384 | Geographical features |
| 74 | DPCP2_Tilt (RNA)_pos10 | iFeatures | 162 | Region-based geo 392 | Geographical features |
| 75 | DPCP2_Shift (RNA)_pos11 | iFeatures | 163 | Region-based geo 394 | Geographical features |
| 76 | DPCP2_Slide (RNA)_pos11 | iFeatures | 164 | Region-based geo 402 | Geographical features |
| 77 | DPCP2_Twist (RNA)_pos30 | iFeatures | 165 | Region-based geo 404 | Geographical features |
| 78 | DPCP2_Rise (RNA)_pos37 | iFeatures | 166 | Region-based geo 412 | Geographical features |
| 79 | DPCP2_Roll (RNA)_pos37 | iFeatures | 167 | Region-based geo 414 | Geographical features |
| 80 | DPCP2_Rise (RNA)_pos38 | iFeatures | 168 | Region-based geo 422 | Geographical features |
| 81 | 100-way PhyloP 1 | Conservation scores | 169 | Region-based geo 424 | Geographical features |
| 82 | 100-way PhyloP 2 | Conservation scores | 170 | Region-based geo 432 | Geographical features |
| 83 | 100-way PhyloP 3 | Conservation scores | 171 | Region-based geo 442 | Geographical features |

| | | | | | |
|---|---|---|---|---|---|
| 84 | 100-way PhyloP 4 | Conservation scores | 172 | Region-based geo 444 | Geographical features |
| 85 | 100-way PhyloP 5 | Conservation scores | 173 | Region-based geo 452 | Geographical features |
| 86 | 100-way PhyloP 9 | Conservation scores | 174 | Region-based geo 454 | Geographical features |
| 87 | Distance-based geo 2 | Geographical features | 175 | Region-based geo 501 | Geographical features |
| 88 | Distance-based geo 3 | Geographical features | | | |

**Table S5.** The performance of AI-m6ARS trained with 6,396 features under different classifiers under 5-fold cross-validation.

| **Algorithm** | **Area Under the Curve** | **Balanced Accuracy** | **F1-score** | **Matthews Correlation Coefficient** |
|---|---|---|---|---|
| **Light Gradient Boosting Machine (LightGBM)** | **0.869 ± 0.005** | **0.594 ± 0.004** | **0.303 ± 0.010** | **0.316 ± 0.008** |
| Gradient Boosting (GB) | 0.867 ± 0.007 | 0.588 ± 0.003 | 0.288 ± 0.008 | 0.306 ± 0.006 |
| Explainable Boosting Machine (EBM) | 0.858 ± 0.008 | 0.589 ± 0.005 | 0.291 ± 0.014 | 0.309 ± 0.011 |
| Extreme Gradient Boosting (XGBoost) | 0.856 ± 0.005 | 0.593 ± 0.006 | 0.298 ± 0.015 | 0.299 ± 0.017 |
| Adaptive Boosting (ADABoost) | 0.848 ± 0.007 | 0.605 ± 0.005 | 0.323 ± 0.009 | 0.309 ± 0.004 |
| Random Forest (RF) | 0.824 ± 0.006 | 0.502 ± 0.000 | 0.006 ± 0.001 | 0.045 ± 0.006 |

| | | | | |
|---|---|---|---|---|
| Extra Trees | 0.794 ± 0.006 | 0.502 ± 0.001 | 0.010 ± 0.004 | 0.053 ± 0.016 |
| Multilayer perceptron (MLP) | 0.684 ± 0.014 | 0.511 ± 0.009 | 0.047 ± 0.037 | 0.077 ± 0.045 |
| K-Nearest Neighbors (KNN) | 0.681 ± 0.005 | 0.579 ± 0.006 | 0.249 ± 0.013 | 0.192 ± 0.013 |
| LogisticRegression | 0.548 ± 0.023 | 0.500 ± 0.000 | 0.002 ± 0.002 | 0.003 ± 0.006 |

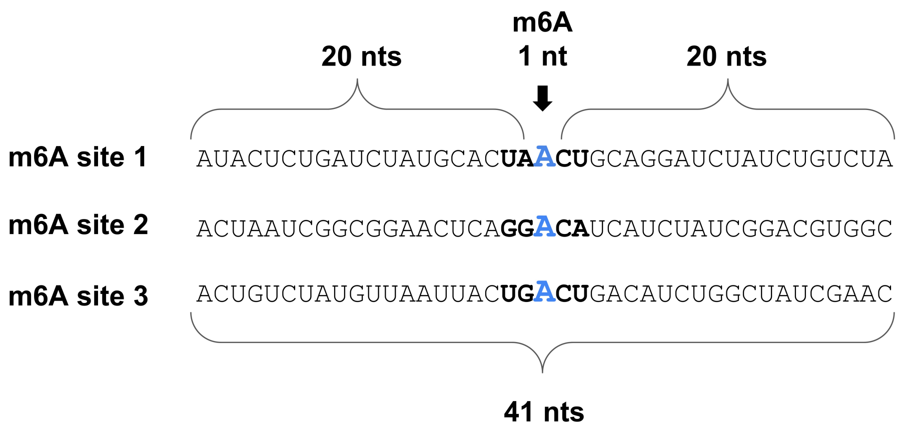

**Figure S1. Graphical representation of experimentally validated m6A sites generated from the miCLIP experiment.** 41-nucleotide sequences, centred on the m6A site (highlighted in blue), were used as RNA sequence representation for validated functional and non-functional m6A sites. The consensus DRACH motifs are highlighted in bold.

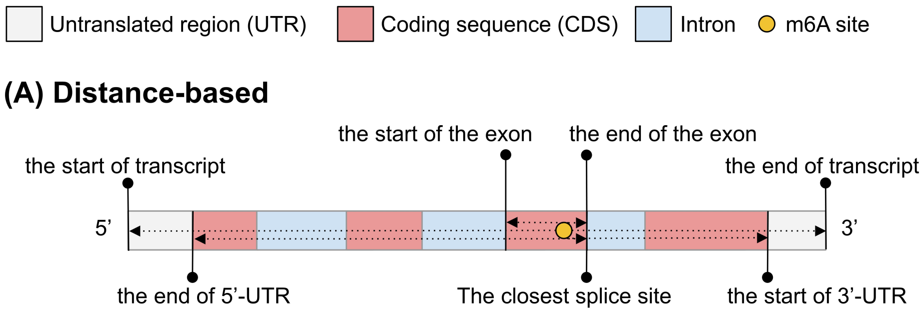

#### (B) Fragment-based

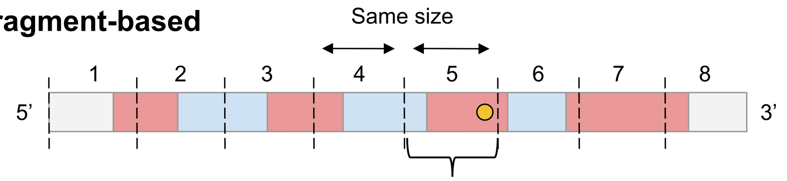

| Number of fragment | Length of fragment | Index of fragment | m6A | Exon | 5’-UTR | CDS | Intron | 3’-UTR |
|---|---|---|---|---|---|---|---|---|
| 8 | 100 | 5 | 1 | 0.8 | 0 | 0.8 | 0.2 | 0 |

#### (C) Region-based

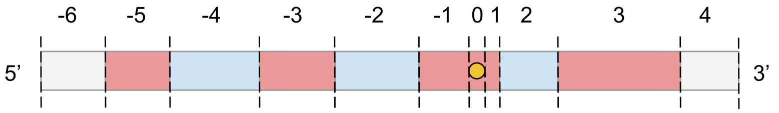

| Region index | -25 | ... | -2 | -1 | 0 | 1 | 2 | ... | 25 |
|---|---|---|---|---|---|---|---|---|---|
| Region length | 0 | ... | 60 | 30 | 1 | 20 | 50 | ... | 0 |
| Proportion | -1 | ... | 0.08 | 0.04 | 0.00 | 0.02 | 0.06 | ... | -1 |
| $-\log_2$(Proportion) | -1 | ... | 3.74 | 4.74 | 9.64 | 5.32 | 4.00 | ... | -1 |
| m6A | 0 | ... | 0 | 0 | 1 | 0 | 0 | ... | 0 |
| 5’-UTR | 0 | ... | 0 | 0 | 0 | 0 | 0 | ... | 0 |
| CDS | 0 | ... | 0 | 1 | 1 | 1 | 0 | ... | 0 |
| Intron | 0 | ... | 1 | 0 | 0 | 0 | 1 | ... | 0 |
| 3’-UTR | 0 | ... | 0 | 0 | 0 | 0 | 0 | ... | 0 |
| Exon | 0 | ... | 0 | 1 | 1 | 1 | 0 | ... | 0 |

**Figure S2. Geographical features implemented in AI-m6ARS. AI-m6ARS encodes three types of geographical features: Distance, Fragment, and Region.** (A) Distance-based features capture relative positional information, including the type of region where m6A resides and distances to key geographical landmarks within a transcript, such as transcript boundaries, exons, splice sites, 5′-UTR, and 3′-UTR. (B) Fragment-based features divide transcripts into uniform-size fragments, with nine features characterising each fragment based on length and composition for each genomic region. (C) Region-based features partition transcript into regions centred on the target site, with ten features encoding m6A site presence, region length, and region type for each region.

#### General Information

| Provided ID | Transcript ID | Gene ID | Chromosome | Length |
|---|---|---|---|---|
| ENST00000005178 | ENST00000005178 | ENSG00000004799 | 7 | 3601 |

| Start | Stop | Strand | Transcript type | No. predicted m6A sites |
|---|---|---|---|---|
| 95583499 | 95596516 | - | mRNA | 1 |

#### Prediction result of m6A modification sites

| m6A position (transcript) | m6A position (genome) | Prediction | Probability | DRACH motif |
|---|---|---|---|---|
| 151 | 95596366 | No | 0.228 | G A A C C |
| 1434 | 95585666 | No | 0.414 | A A A C A |
| 3358 | 95583742 | Yes | 0.502 | A G A C A |

#### The distribution of m6A modification sites : ENST00000005178

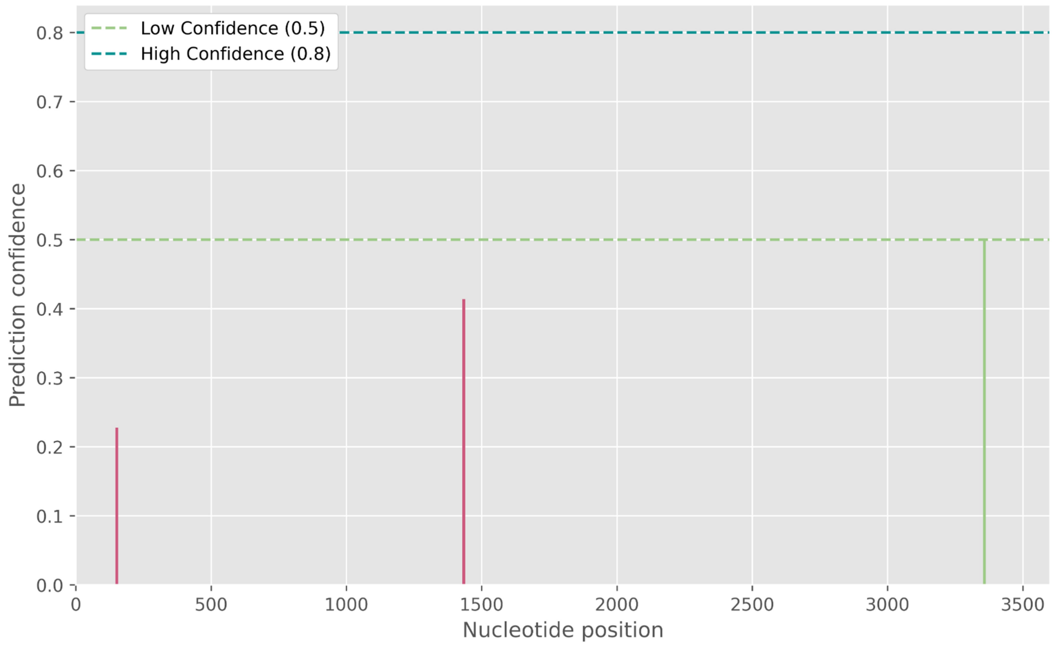

**Figure S3. Detail page of AI-m6ARS web server.** The detail page is accessible through the "View Details" button on the main result page (refer to Figure 4B). The page is structured into three sections: General Information, Prediction Results, and Distribution of Modification Sites. The first section presents comprehensive information regarding transcripts, including Ensembl IDs, chromosomal positions, transcript types, and the number of predicted m6A sites. The second section exhibits tabular information on m6A positions, accompanied by their respective predictions, probabilities, and DRACH motif sequences. The final section presents the distribution graph of m6A modification sites, providing position-specific information on the predicted modifications.
